# Impact of Data Quality on Deep Learning Prediction of Spatial Transcriptomics from Histology Images

**DOI:** 10.1101/2025.09.04.674228

**Authors:** Caleb Hallinan, Calixto-Hope G. Lucas, Jean Fan

## Abstract

Spatial transcriptomic technologies enable high-throughput quantification of gene expression at specific locations across tissue sections, facilitating insights into the spatial organization of biological processes. However, high costs associated with these technologies have motivated the development of deep learning methods to predict spatial gene expression from inexpensive hematoxylin and eosin-stained histology images. While most efforts have focused on modifying model architectures to boost predictive performance, the influence of training data quality remains largely unexplored. Here, we investigate how variation in molecular and image data quality stemming from differences in spatial transcriptomic technologies impact deep learning-based gene expression prediction from histology images. To identify the aspects of data quality that impact predictive performance, we conducted *in silico* ablation experiments, which showed that increased sparsity and noise in molecular data degraded predictive performance, while *in silico* rescue experiments via imputation provided only limited improvements that failed to generalize beyond the test set. Likewise, reduced image resolution can degrade predictive performance and further impacts model interpretability. We further demonstrate that these data quality-driven effects are reproducible across multiple spatial transcriptomics datasets and remain consistent when using alternative feature extractors and model architectures. Overall, our results show how improving data quality provides an orthogonal strategy to tuning model architecture in spatial transcriptomics-based predictive modeling, highlighting the need to account for technology-specific limitations that directly impact data quality when developing predictive methodologies.

## Introduction

Spatial transcriptomic (ST) technologies enable the measurement of gene expression with spatial context across tissue sections with varying molecular resolution and throughput. These spatial molecular measurements are often accompanied by a hematoxylin and eosin (H&E)-stained image of the tissue section used for the ST assay. Although ST experiments can cost several thousand dollars per sample, H&E-stained images are relatively inexpensive and routine in clinical and research histopathology. By leveraging such matched H&E-stained images and gene expression data, machine learning models can be trained to predict molecular information directly from histology images. This greatly extends the utility of both archived and newly generated histology images for molecular inference, potentially bypassing the need for expensive ST technologies.

The quality of molecular data can vary widely across ST technologies, reflecting the unique technical constraints inherent to each technology. Imaging-based technologies (for example, STARmap [1], MERFISH [2], NanoString CosMx, and 10x Xenium) provide sub-cellular resolution and typically higher sensitivity but are currently limited to smaller gene panels and can suffer from optical signal overlap or off-target probe-binding issues [3, 4]. In contrast, sequencing-based technologies (for example, Slide-seqV2 [5], DBiT-seq [6], and 10x Visium) provide transcriptome-wide coverage but at generally lower spatial resolution and can be prone to lateral diffusion between spots, problematic spots, as well as PCR amplification biases and drop-outs that yield sparse, noisy counts [7–10]. In addition to these ST technology-specific limitations impacting the molecular data quality, the quality of histology images can also vary through differences in staining protocols, scanner optics, image compression, and other factors that alter color balance, contrast, and effective pixel size across datasets [11, 12]. Although multiple sources contribute to variation in data quality, their collective impact on the performance and interpretability of downstream predictive models remains largely unexplored.

A growing body of work has applied deep learning models, varying in terms of the convolutional, graph, or transformer-based architectures used, to predict gene expression from histology images using ST data for training [13, 14]. At first, these approaches were almost exclusively trained and evaluated on data from sequencing-based technologies, such as Visium, where predictive performance, as measured by the Pearson correlation coefficient between predicted and observed gene expression, often averaged around 0.3. More recent studies have begun training and evaluating models using data from imaging-based technologies, such as Xenium, reporting a higher predictive performance [15, 16]. However, prediction performance remains highly variable across datasets, tissue types, and models, highlighting the need for further investigation into factors that impact prediction performance [17].

In light of this, our study assesses the impact of training data quality on the performance of gene expression prediction from histology images using deep learning models. To this end, we first utilized two previously published breast cancer datasets from serial tissue sections assayed by a sequencing-based technology (Visium) and an imaging-based technology (Xenium). The Visium data is accompanied by the 10x Genomics provided high-resolution histology image and a re-scanned whole-slide H&E-stained image (WSI), whereas the Xenium data includes a high-resolution WSI of the same tissue section [18]. We find that when using Xenium data as the input training data compared to when using Visium data, the average prediction performance across genes increased. To determine what specific data quality factors in the input training data may contribute to the observed prediction performance differences between technologies, we conducted a series of *in silico* ablation experiments. First, we performed molecular ablation experiments by introducing expression sparsity and noise in the higher-performing Xenium data, which recapitulated the the lower-performance observed when training on Visium data. In addition, molecular rescue experiments aimed at enhancing the molecular data quality of the Visium data through gene expression imputation resulted in overfitting and poor generalizability, indicating that such artificial augmentation may not substitute higher quality data. We next examined the role of image quality by reducing the resolution of the Xenium histology image, which led to corresponding decreases in predictive performance. In contrast, training with the re-scanned Visium whole-slide image yielded modest performance improvements, further highlighting the role of image resolution on prediction performance. To evaluate whether these findings were robust to model choice, we assessed two alternative approaches: (1) replacing the ResNet50 feature extractor with the UNI histology foundational model and (2) implementing the spatial-transcriptomics specific model RedeHist, finding consistent relative performance trends. Furthermore, to examine the generalizability of data quality effects across tissue and technologies, we extended our framework to a colon adenocarcinoma dataset comprising of Xenium 5K, VisiumHD, and CosMx 6K, where similar technology-specific data quality impacts were observed. Overall, our results underscore the critical role of training data quality in spatial gene expression prediction and demonstrates that training on higher quality data can improve prediction performance.

## Results

### Data quality benchmarking on paired Visium and Xenium spatial transcriptomics datasets

To investigate how ST training data quality affects gene expression prediction from histology images via deep learning, we first leveraged serial section breast cancer datasets generated by Visium CytAssist and Xenium (Figure 1A) [18]. Because a single tissue section cannot undergo two distinct ST assays, serial sections allow us to control for biological variability and focus on technical variability across the different ST technologies. Still, slight differences in the region of the tissues profiled as well as image and molecular data resolution persist. Therefore, to better ensure that differences in gene expression prediction performance can be attributed to the training regime rather than to confounding variation between the training datasets, we standardize as many aspects of the input as possible. To this end, we first aligned the Visium histology image to the Xenium histology image using STalign, bringing both datasets into a common co-ordinate system (Figure 1B) [19]. This enabled the molecular data from either technology to be visualized and utilized with either histology image. We then identified the shared region between the two datasets and rasterized the gene expression to a common spatial resolution (Figure 1C), as demonstrated by Aihara et al. [20]. By minimizing such sample-specific differences, this preprocessing enables controlled comparisons across all combinations of molecular and imaging data. Using this setup, we extracted histology image patches that correspond to the rasterized molecular data and trained identical deep learning models composed of a pretrained ResNet50 feature extractor, a four-layer multilayer perceptron, and a final linear output layer to predict gene expression at the patch level for the 306 genes common to both ST assays (Figure 1D). We evaluated model prediction performance by computing Pearson correlation coefficients (PCC) and range-normalized root mean square error (rMSE) between true and predicted patch-level gene expression on a held-out test set (15% of patches) and, in some cases, on an independent serial section Xenium replicate dataset (Rep2) to further assess generalizability. To determine which factors of data quality influence prediction performance and interpretability, we performed a series of *in silico* ablation experiments including increasing expression sparsity and noise, applying Gaussian blur to simulate lower-resolution images, as well as gene expression imputation in an attempt to remove sparsity and noise, and then reassessed prediction performance as well as Grad-CAM saliency (Figure 1E).

**Figure 1:**
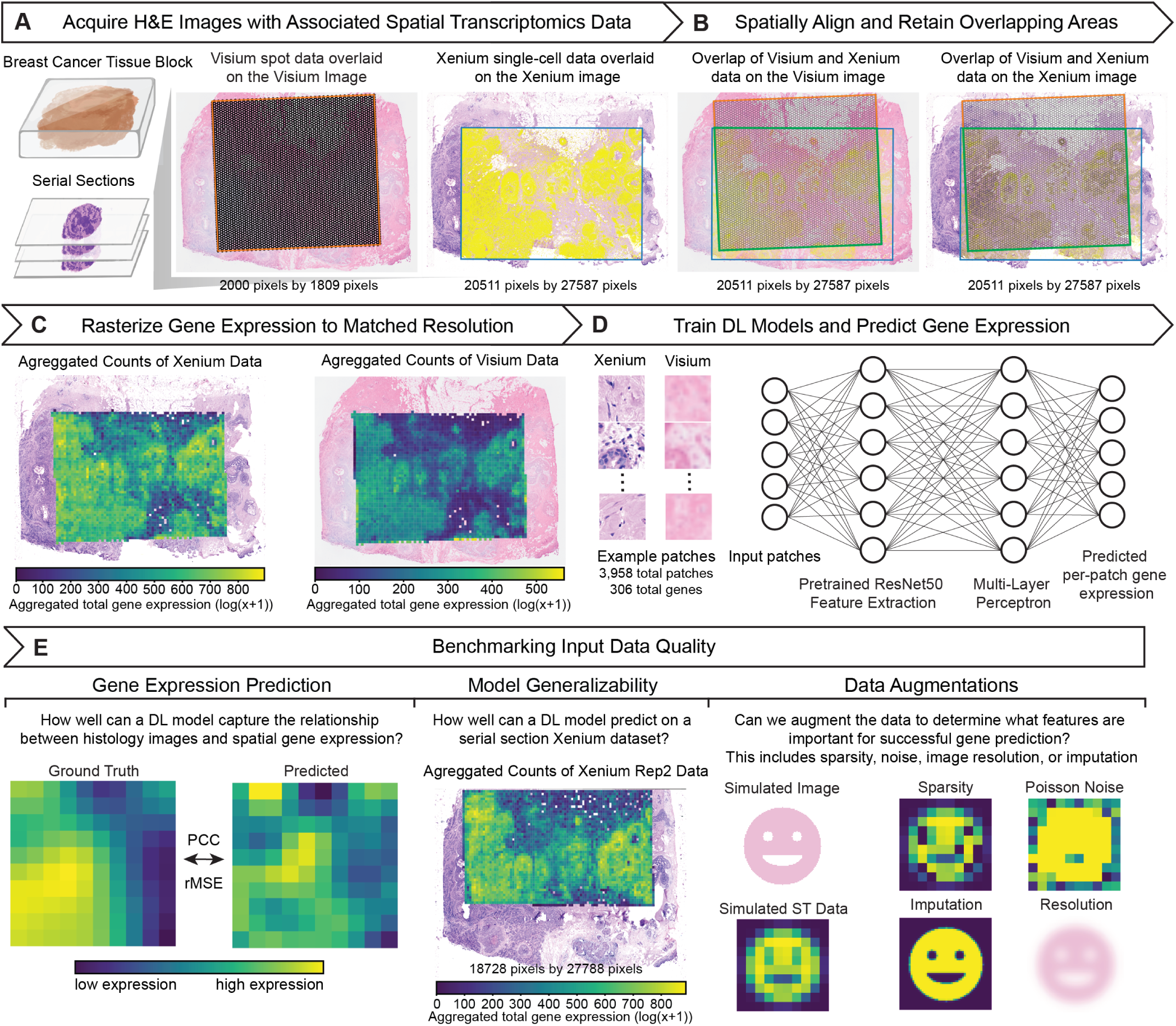
Overview of the data-quality benchmarking pipeline. **A.** Acquisition of paired breast cancer spatial transcriptomics datasets and histology images from 10x Visium and Xenium. **B.** Co-registration of Visium and Xenium histology slides into a common coordinate system. The green box highlights the overlapping region retained between the two technologies. **C.** Rasterization of gene counts onto a uniform grid matched to Visium spot resolution, followed by extraction of the overlapping tissue region. Expression is visualized as patches. **D.** Training of deep learning models to predict per-patch gene expression from histology image patches. **E.** Performance evaluation on held-out replicates, comparison across technologies, and ablation experiments of inputs.

Applying this approach, we directly compared the predictive performance of our deep learning model when trained on each technologies’ associated histology image and corresponding molecular data (Figure 2). For every gene, we computed the PCC between the ground-truth expression values and model predictions on the held-out test set, using five independently trained models to account for training variability. We found that the PCC distribution across genes is notably shifted toward higher values, indicative of better prediction performance, when trained on Xenium data compared to Visium data (mean PCC = 0.715 for Xenium vs. 0.519 for Visium; Figure 2A), representing an increased average prediction performance across genes of approximately 38%. Likewise, in the gene-by-gene scatterplot (Figure 2B), almost all points lie above the diagonal indicating that prediction performance when training on Xenium data was consistently higher than with Visium data for almost all genes. Evaluating the complete distribution of ground-truth and predicted expression values for held-out test set patches reveals that the PCC is not driven by outliers (Supplementary Figure 1). To further ensure this trend holds across metrics, we also examined the range-normalized root mean square error (rMSE) and found that gene expression predictions compared to the ground-truth expression values exhibited higher normalized rMSE, indicative of worse performance, when trained on Visium data compared to Xenium data (Supplementary Figure 2). Plots of observed versus predicted expression for four representative genes (*HDC*, *ANKRD30A*, *AHSP*, *GZMK*) highlight varying performance outcomes: Xenium-trained models outperform Visium for *HDC*, both models perform similarly well for *ANKRD30A* and poorly for *AHSP*, while Visium-trained models slightly outperforms Xenium for *GZMK* (Figure 2C).

**Figure 2:**
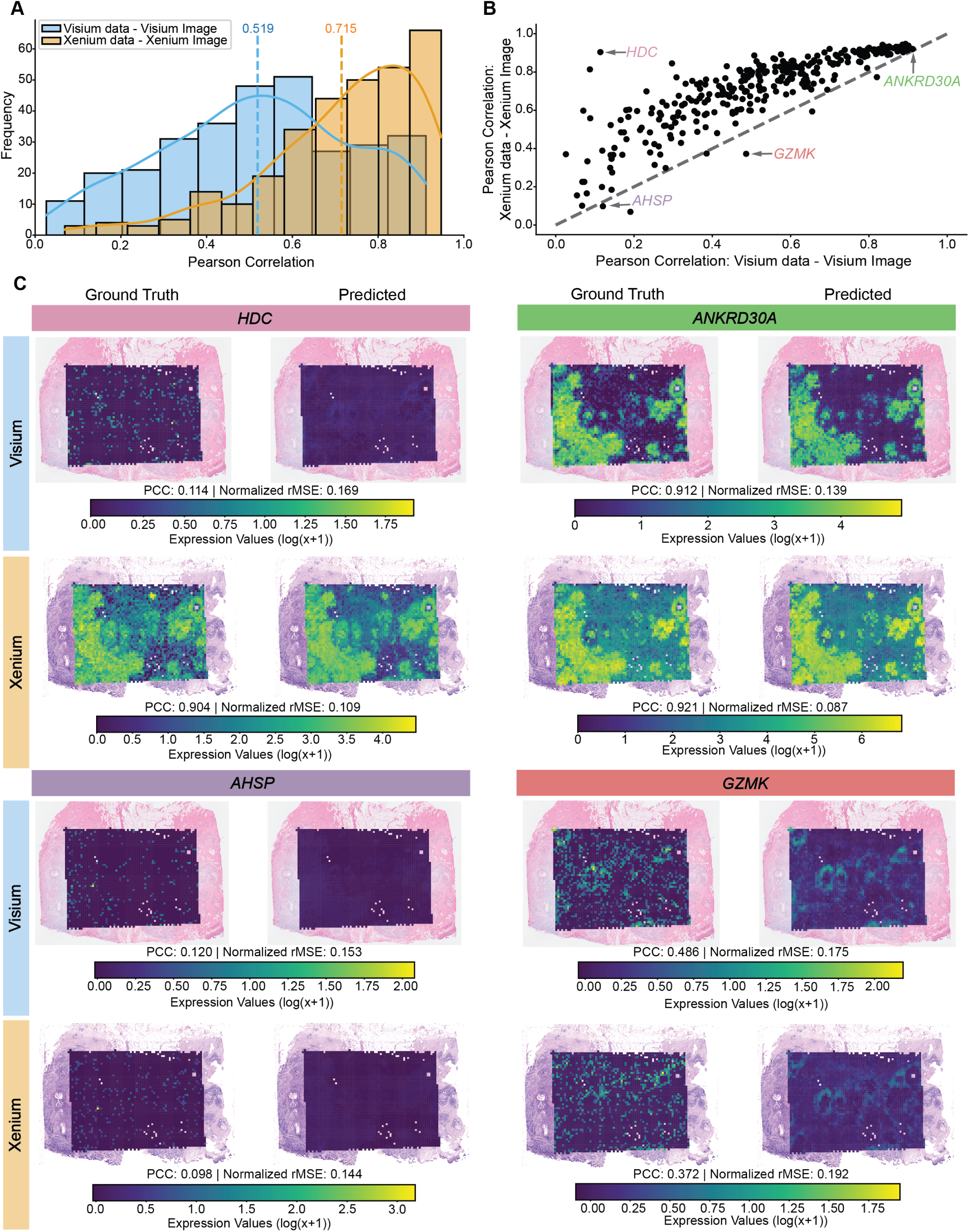
Spatial gene expression prediction comparison using Visium vs.Xenium data. **A.** Histogram showing the distribution of Pearson correlation coefficients for gene expression predictions using Visium and Xenium data. The dotted vertical line denotes the mean PCC, and the solid curved line traces the density estimate. Results are computed on the held-out test set and represent the average performance across five independently trained models. **B.** Scatterplot comparing the Pearson correlation coefficients of predictions from Visium and Xenium data. The gray dotted line denotes x=y, and select genes corresponding to (C) labeled. **C.** Representative examples of ground truth and predicted gene expression for *HDC*, *ANKRD30A*, *AHSP*, and *GZMK* in both the Visium and Xenium datasets. Predicted gene expressions are visualized for the full dataset, while the performance metrics (PCC and normalized rMSE) are computed from the held-out test set only.

To confirm these findings were robust to model choice, we evaluated two additional architectures. The first used the same overall structure shown in Figure 1D but replaced the feature extractor with the UNI histology foundational model [21], while the second used RedeHist [22], a model specifically developed for gene expression prediction from histology images using spatial transcriptomics data. Using UNI resulted in a slight increase in PCC for both Xenium and Visium results (mean PCC = 0.734 for Xenium vs. 0.529 for Visium; Supplementary Figure 3A-B) and decrease in normalized rMSE, but the overall trend in prediction performance when trained on Xenium data compared to Visium data remained the same. Using RedeHist, there was a modest decrease in PCC and increase in normalized rMSE for both Xenium and Visium data (mean PCC = 0.633 for Xenium vs. 0.385 for Visium; Supplementary Figure 3C-D), but again the overall trend remained the same.

Overall, these results indicate that models trained on Xenium’s paired molecular data and histology image achieve better gene expression predictive performance than those trained on the Visium molecular data and its corresponding histology image. Whether this advantage arises primarily from differences in molecular data quality or from image quality or both remains to be determined in our subsequent analyses.

### Molecular data quality can impact spatial gene expression prediction performance

To disentangle the impact of molecular data quality from image data quality, we next held the image data constant while swapping molecular data between technologies. We leveraged the high-resolution histology image from the Visium dataset and trained two identical models: one on Visium molecular data, the other on Xenium molecular data. Under these conditions, Xenium molecular data outperformed Visium molecular data (mean PCC = 0.605 vs. 0.519; Figure 3A), and most genes exhibited higher PCCs when trained with the Xenium molecular data (Figure 3B). We next repeated this with the WSI from the Xenium dataset. Again, models trained using Xenium molecular data outperformed Visium molecular data (mean PCC = 0.715 for Xenium vs. 0.492 for Visium; Figure 3C-D). These trends further held for normalized rMSE (Supplementary Figure 4). This demonstrates that, irrespective of the image data, the molecular data used for training has an effect on gene expression prediction performance.

**Figure 3:**
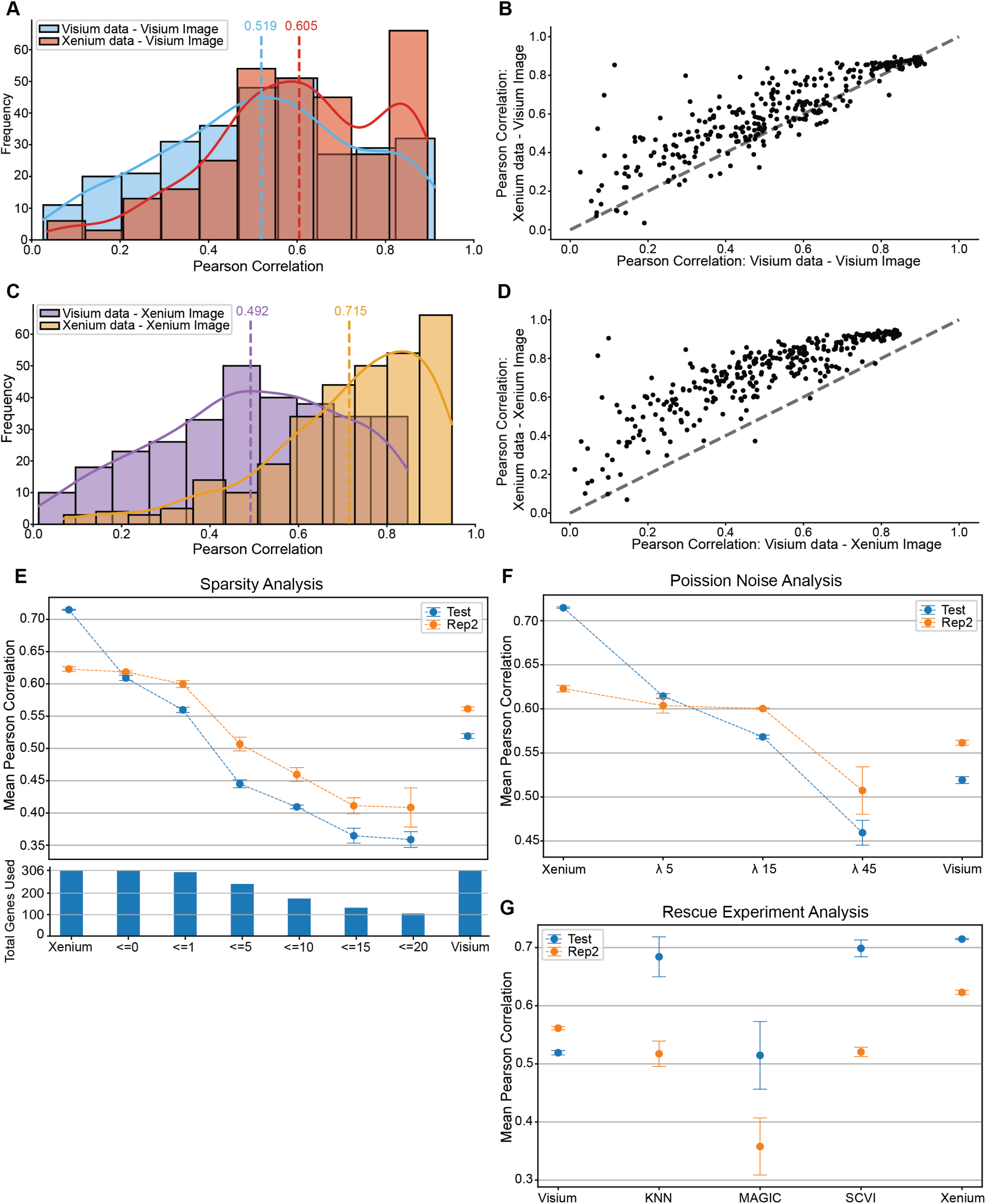
Impact of molecular data quality on spatial gene expression prediction. **A.** Histogram of Pearson correlation coefficients for gene expression predictions using Visium and Xenium data with the Visium image. The dotted vertical line denotes the mean PCC, and the solid curved line traces the density estimate. **B.** Scatterplot comparing PCC values from Visium and Xenium data with the Visium image on the test set, averaged across five models. The gray dotted line denotes x=y. **C.** Histogram of PCC values for predictions using Visium and Xenium data with the Xenium image. The dotted vertical line denotes the mean PCC, and the solid curved line traces the density estimate. **D.** Scatterplot comparing PCC values from Visium and Xenium data with the Xenium image on the test set, averaged across five models. The gray dotted line denotes x=y. **E.** Scatterplot comparing PCC values between Xenium, an increasing amount of sparsity in the Xenium dataset, and the Visium results on the test and replicate 2 Xenium data. The dotted line indicates the dataset used, and error bars represent the standard error across five runs. The histogram below denotes the total number of genes used to calculate the mean PCC. **F.** Scatterplot comparing PCC values between Xenium, an increasing amount of Poisson noise in the Xenium dataset, and the Visium results on the test and replicate 2 Xenium data. The dotted line indicates the dataset used, and error bars represent the standard error across five runs. **G.** Scatterplot comparing PCC values between Visium, various imputation methods on the Visium dataset, and the Xenium results on the test and replicate 2 Xenium data. The dotted line indicates the dataset used, and error bars represent the standard error across five runs.

To pinpoint what molecular data quality factors may drive these performance differences, we next performed a series of *in silico* ablation experiments in which we systematically decrease molecular data quality and evaluate its impact on prediction performance. First, we examined expression sparsity, defined here as the proportion of zero counts across patches for each gene. Consistent with previous research, we note that sparsity is more pronounced in the molecular data from Visium than Xenium (Supplementary Figure 5). Having confirmed Xenium data’s better performance, we increased sparsity in the patch-resolution Xenium gene expression count matrix to match the sparsity in the Visium gene expression count matrix. To do this, we compared corresponding patches from the Visium and Xenium datasets, and wherever the Visium count matrix exhibited a value of ≤ 0, we set the corresponding patch in the patch-resolution Xenium count matrix value to 0. We repeated this process for Visium expression thresholds of ≤ 1, ≤ 5, ≤ 10, ≤ 15, and ≤ 20. As sparsity increased, mean PCC on the held-out test set steadily declined (Figure 3E). To ensure this effect was not an artifact of the held-out test data and to access model generalization on a new dataset, we repeated the evaluation on an independent Xenium serial section (Rep2). Here too, we observed a similar downward trend in PCC as sparsity increased (Figure 3E). Notably, training with Xenium molecular data with increased sparsity using a ≤ 1 threshold produced comparable performance to when training with the Visium molecular data. Next, we examined expression noise, defined as random fluctuations in gene counts around their true values. To simulate greater expression noise, we added Poisson distributed noise into the patch-resolution Xenium count matrix across a series of rate parameters (*λ*). As we increased *λ* to simulate more expression noise, mean PCC on the held-out test set steadily declined (Figure 3F). We again repeated the evaluation on the independent Xenium serial section (Rep2) to confirm this trend. Overall, these results suggest molecular data sparsity and noise as potential drivers of decreased prediction performance.

Finally, given that increased sparsity and noise can degrade prediction performance, we asked whether we could rescue the Visium molecular data via gene imputation. Gene imputation refers to techniques that infer and fill in missing or zeroed expression values by borrowing information from similar cells or genes across the dataset or smoothing expression profiles to mitigate sparsity and noise issues. We applied three representative imputation methods: KNN [23], MAGIC [24], and SCVI [25] to the Visium molecular data, retrained our models, and evaluated PCC on the test set and on the independent replicate (Figure 3G). Applying these imputation methods boosted the PCC when evaluated on the held-out test set. However, when we repeated evaluation on the independent Xenium replicate, applying these imputation methods led to a consistent decrease in the PCC. This suggests that while imputation can mask sparsity and noise to aid model training, it may introduce biases that limit robustness and generalizability to new samples.

### Image data quality can impact spatial gene expression prediction performance

To isolate the effect of image data quality from molecular data quality, we held the molecular input constant and swapped histology images between technologies. First, using Visium molecular data, we trained models on both the Visium high-resolution image (2000px by 1809px) and the higher resolution Xenium WSI (20511px by 27587px). The resulting PCC distributions were comparable (mean PCC = 0.519 with Visium image vs. 0.492 with Xenium image; Figure 4A). Gene-wise comparisons of PCC remained along the diagonal (Figure 4B), and normalized rMSE values were similarly low (Supplementary Figure 6A), indicating that image resolution had little effect on prediction performance. Next, using Xenium molecular data, we repeated the experiment and observed a larger performance gap (mean PCC = 0.605 with Visium image vs. 0.715 with Xenium image; Figure 4C). Gene-by-gene scatterplots (Figure 4D) and normalized rMSE comparisons (Supplementary Figure 6B) demonstrate consistent trends. As such, training on higher resolution histology images can improve prediction performance depending on the quality of the corresponding molecular data. Consistent with this, when using the Visium WSI (20511px by 27587px) instead of the CytAssist associated high-resolution image, this trend persisted, with higher PCCs for models trained on the Xenium molecular data compared to the Visium molecular data (mean PCC = 0.678 for Xenium vs. 0.526 for Visium), accompanied by a decrease in normalized rMSE for the Xenium-trained models and slight increase for the Visium-trained models (Supplementary Figure 7A-B). Likewise, models trained on the Visium data showed a slight increase in PCC (from 0.519 to 0.526), accompanied by a slight increase in normalized rMSE (from 0.179 to 0.181; Supplementary Figure 7C-D).

**Figure 4:**
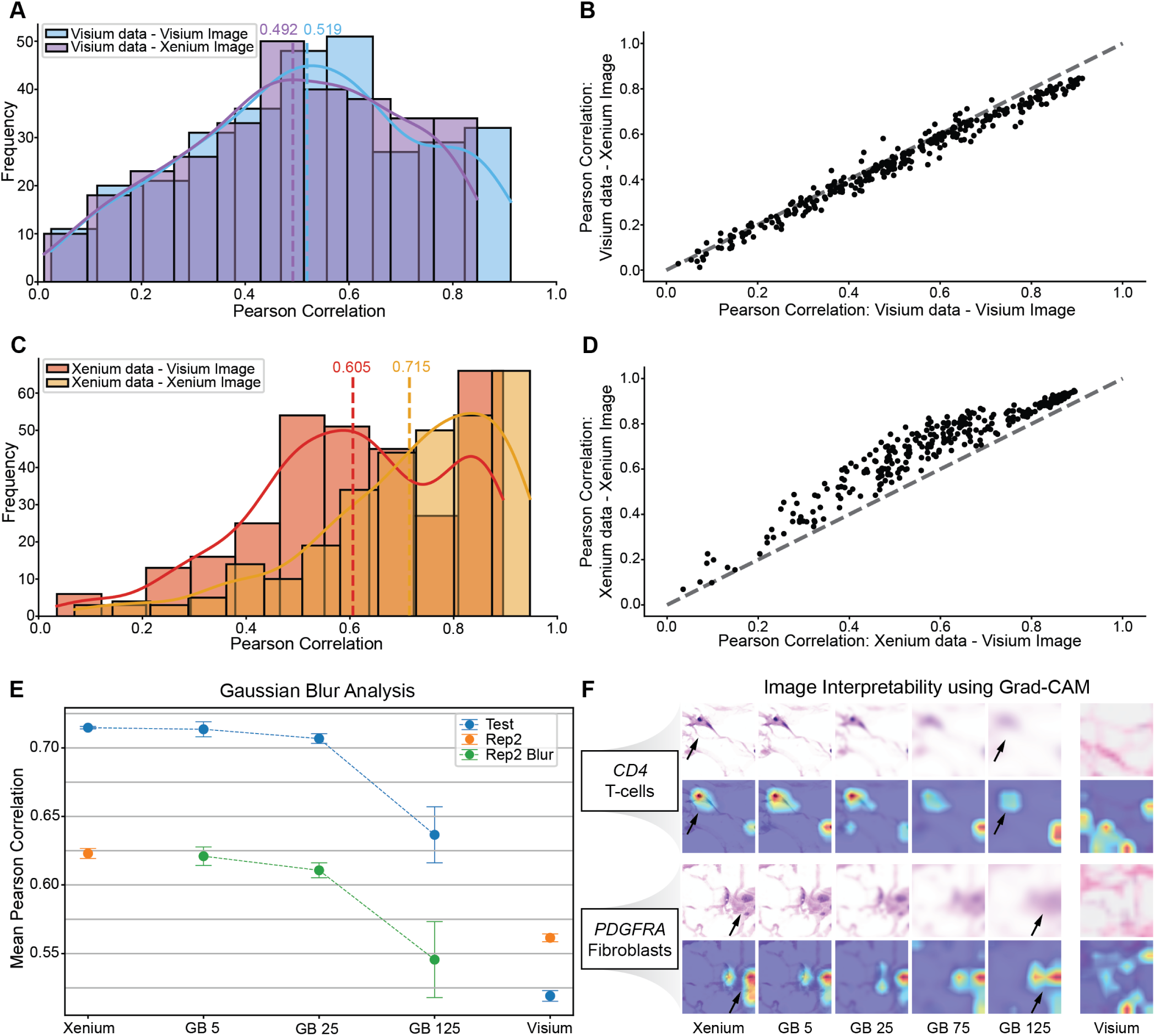
Impact of imaging data quality on spatial gene expression prediction. **A.** Histogram showing the distribution of Pearson correlation coefficients for gene expression predictions using Visium data with the Visium and Xenium images. The dotted vertical line denotes the mean PCC, and the solid curved line traces the density estimate. Results are computed on the test set and represent the average performance across five independently trained models. **B.** Scatterplot comparing the Pearson correlation coefficients of predictions from Visium data with the Visium and Xenium images, based on the test set and averaged over five models. The gray dotted line denotes x=y. **C.** Histogram showing the distribution of Pearson correlation coefficients for gene expression predictions using the Xenium data with the Visium and Xenium image. The dotted vertical line denotes the mean PCC, and the solid curved line traces the density estimate. Results are computed on the test set and represent the average performance across five independently trained models. **D.** Scatterplot comparing the Pearson correlation coefficients of predictions from Xenium data with the Visium and Xenium image, based on the test set and averaged over five models. The gray dotted line denotes x=y. **E.** Scatterplot of mean Pearson correlation coefficients on both the test set and the Replicate 2 Xenium section, comparing the Xenium, Xenium images with increasing Gaussian blur, and Visium results (all applied with the same blur levels). The dotted line indicates the dataset used, and error bars represent the standard error of the mean across five independent model runs. **F.** Grad-CAM heatmaps for two select genes: *CD4* (T-cell marker) and *PDGFRA* (fibroblast marker).

To pinpoint what imaging data quality factors may drive these performance differences, we simulated lower-resolution images by applying Gaussian blur with kernels of size 5, 25, and 125 to the Xenium histology images. Under matched conditions, training and testing on equally blurred images, PCC slowly degraded with decreasing simulated image resolution (Figure 4E). These results suggest that image resolution has a measurable impact on gene expression prediction performance, with progressively blurred images leading to decreased prediction performance.

Finally, beyond the impact of image resolution on gene expression prediction performance, we sought to assess its impact on model interpretability. To this end, we employed Grad-CAM [26], which projects prediction gradients onto convolutional feature maps to generate heatmaps that highlight the image regions most influential for a given output. This technique is especially useful because it can reveal whether the model focuses on potentially biologically meaningful structures, such as cells or nuclei consistent with what histopathologists may focus on, rather than background artifacts. We applied Grad-CAM to predictions for select well-predicted genes with high PCCs including *CD4* (PCC = 0.79), a T-cell marker, and *PDGFRA* (PCC = 0.76), a fibroblast marker (Figure 4F). On high-resolution images, the resulting heatmaps highlighted cellular and nuclear regions. As we introduced increasing levels of blur to simulate reduced resolution, the heatmaps became diffuse and lost alignment with these structural landmarks. These results demonstrate that lower image resolution may not only decrease gene expression prediction performance but also reduce reliable interpretation of the tissue features that drive model predictions.

### Reproducibility of data quality effects across spatial transcriptomics technologies

To assess whether the observed data quality effects could be reproduced with additional spatial transcriptomics technologies and tissues, we next applied our framework to a colon adenocarcinoma (COAD) dataset comprising of serial tissue sections assayed with three spatial transcriptomics technologies: Xenium 5K, VisiumHD, and CosMx 6K [27]. Xenium 5K expands the targeted panel to approximately 5000 genes, compared with fewer than 400 in the original Xenium panel, although this comes with slightly reduced sensitivity [28]. VisiumHD provides higher molecular resolution than Visium CytAssist, enabling measurements at patch sizes as small as 2*µ*m x 2*µ*m, while also eliminating gaps between spots. CosMx 6K is an imaging-based technology analogous to Xenium 5K but uses a modified two-probe chemistry rather than padlock probes [29]. Using the paired H&E-stained whole-slide images and spatial transcriptomic measurements, we rasterized all datasets to a common resolution of 55*µ*m x 55*µ*m patches and aggregated gene expression within each patch. We then trained five independently trained deep-learning models to predict patch-level gene expression from histology images and evaluated model prediction performance using PCC and rMSE on a held-out test set (15% of patches) for each dataset.

Consistent with our previously observed trends, prediction performance was impacted by the training data, as the PCC distribution across genes is shifted toward higher values when trained on Xenium 5K data compared to VisiumHD data (mean PCC = 0.568 for Xenium 5K vs. 0.434 for VisiumHD; Supplementary Figure 8A). Likewise, the normalized rMSE was lower for Xenium 5K when compared to VisiumHD (mean rMSE = 0.148 for Xenium 5K vs. 0.162 for VisiumHD; Supplementary Figure 8B). Gene-wise comparisons of PCC between Xenium 5K and VisiumHD further highlight this difference, with most genes lying above the diagonal in the scatterplot (Supplementary Figure 8C), indicating consistently improved prediction performance when training on Xenium 5K data relative to VisiumHD data. In contrast, models trained on CosMx 6K data displayed a distinct performance distribution. The PCC distribution was largely unimodal, rather than bimodal as observed for VisiumHD and Xenium 5K, with a mean PCC that fell between the two technologies (mean PCC = 0.532) but accompanied by higher overall error (mean rMSE = 0.200; Supplementary Figure 8A-B). Gene-wise comparisons of PCC between CosMx 6K with VisiumHD and Xenium 5K (Supplementary Figures 8D-E) revealed a low correlation, with genes poorly predicted in VisiumHD and Xenium 5K often better predicted using CosMx 6K data, whereas genes that were well predicted in VisiumHD and Xenium 5K tended to have a worse prediction performance when trained on CosMx 6K data.

To investigate the potential sources of these differences, we examined both image and molecular data quality across technologies. Among the three technologies, the CosMx 6K histology image exhibited the lowest quality based on pathologic assessment, characterized by a pale eosin stain and diffuse hematoxylin staining of nuclei, particularly within epithelial regions (Supplementary Figure 9A). At the molecular level, genes such as *PDYN*, which was better predicted in CosMx 6K relative to Xenium 5K and VisiumHD, showed a spatial pattern that was different from the similar patterns observed between Xenium 5K and VisiumHD, suggesting increased measurement noise (Supplementary Figure 9B). In contrast, genes such as *MAP4*, which was more robustly detected and similarly patterned in Xenium 5K and VisiumHD, was notably sparser in CosMx 6K, corresponding to reduced prediction performance for these genes (Supplementary Figure 9C). This result is not unexpected given prior reports of technology-specific limitations in CosMx data quality and sensitivity [27]. Overall, these results suggest generally that technology-specific data quality features with respect to histology image quality, molecular expression sparsity, and molecular noise can impact which genes are reliably predicted from histology images.

## Discussion

In this study, we evaluated how data quality impacts deep learning–based gene expression prediction from histology images. We leveraged molecular data paired with H&E-stained images from multiple spatial transcriptomic technologies across two tissue types, representing a variety of data quality features. We trained identical deep learning models to predict patch-level gene expression and observed that training on Xenium and Xenium 5K data can achieve quantifiably better gene expression prediction performance than training on Visium, VisiumHD, or CosMx 6K data, with an average prediction performance across genes increase up to 38%. These relative performance trends were consistent across alternative feature extractors and model architectures. Using the breast cancer dataset to isolate the impact of molecular data quality, we held the histology image constant and found that Xenium molecular data outperformed Visium molecular data regardless of which image was used. In molecular ablation experiments, adding noise and sparsity to Xenium molecular data steadily degraded prediction performance. Conversely, in molecular rescue experiments, imputing Visium data boosted performance on the held-out test set but failed to generalize to an independent Xenium replicate. Likewise, to isolate the impact of image data quality, we held the molecular data constant and found Visium molecular data yielded similar prediction performance on both the Visium and Xenium images, whereas Xenium molecular data benefited from the higher-resolution Xenium image. Simulating lower-resolution images by applying Gaussian blur to the Xenium image reduced predictive performance and obscured interpretability via Grad-CAM feature maps. These trends generalized to a colon adenocarcinoma dataset containing Xenium 5K, VisiumHD, and CosMx 6K, where Xenium 5K yielded higher overall prediction accuracy than VisiumHD, while CosMx 6K exhibited gene-specific trade-offs driven by differences in image quality and molecular noise. Overall, these findings underscore the impact of both molecular and imaging data quality in deep learning–based gene expression prediction from histology images. Further, our study presents quantifiable metrics of molecular and imaging data quality in terms of their sparsity, noise, and resolution that can be considered in the curation of training data from ST technologies for developing gene expression prediction methodologies.

Despite these insights, our study has several limitations. Although our comparative analysis demonstrated that training on Visium, VisiumHD, and CosMx 6K data resulted in lower gene expression prediction performance than training on Xenium and Xenium 5K data, potentially attributable to greater sparsity, noise, and other data quality issues in Visium, Xenium and Xenium 5K data can also exhibits their own limitations in data quality. For example, previous research has demonstrated that imaging-based technologies, like Xenium, Xenium 5K, and CosMx, can suffer from off-target probe hybridization, thereby obscuring the target gene’s expression with the off-target gene’s expression [4]. Such artifacts can inflate observed gene expression magnitudes, and because genes with higher mean expression are generally easier to predict, may artificially boost prediction performance when models are trained on such molecular data [15]. We therefore used the Off-Target Probe Tracker (OPT) with a pad-length of 10 to identify 14 of our 306 genes with potential protein-coding off-target genes [4] in the breast cancer data. Removing these genes and retraining our models confirmed that they do not drive our observed trends or affect our conclusions (Supplementary Figure 10). In the case of CosMx, elevated molecular noise or probe-related artifacts may contribute to the apparent performance gains observed for *PDYN* (Supplementary Figure 9B). However, because CosMx probe sequences are not publicly available, we were unable to directly evaluate potential off-target effects in this dataset. Nevertheless, comparing predictions of gene expression using datasets from different ST technologies inherently assumes these different ST technologies are measuring the same underlying genes, which can be obscured by off-target probe binding. Hence, future work is still needed to quantify off-target rates as well as develop robust correction strategies to ensure fair, accurate comparisons across ST technologies.

Similarly for image data quality, beyond image resolution, other features in H&E staining and slide preparation can impact the performance of deep learning models. In particular, H&E staining for Xenium is acquired after the *in situ* gene expression profiling is complete, whereas H&E staining for Visium is acquired before transcriptomic capture. How such protocol variation impacts the quality and generalizability of the H&E-stained images for such predictive modeling purposes remains unclear. Beyond protocol variation, differences in hematoxylin versus eosin concentration, incubation time, and scanner settings can alter color balance, contrast, and texture cues that deep learning models rely on. Common slide artifacts such as air bubbles trapped under the coverslip, uneven tissue folds, tears or wrinkles in the section, dust or debris on the slide and more can further create spurious patterns. Previous studies have shown that these inconsistencies may cause models to learn batch or slide specific artifacts rather than relevant tissue or cellular morphological features [30–33]. As such, we anticipate such common slide artifacts that decrease the quality of the histology image would also impact prediction performance. Likewise, previous studies have sought to address these issues by applying stain-normalization algorithms, color augmentation pipelines, and domain-adaptive foundation models to standardize image appearance across batches and technologies [11, 16, 34, 35]. Future work is needed to evaluate how much such image artifacts impact prediction performance and whether such previously developed strategies are sufficient to rescue poor quality images without introducing biases that limit generalizability. In cases where poor image quality substantially limits predictive performance, re-scanning slides with optimized scanner settings and higher objective lenses could be a practical approach to obtain higher-quality histology images for model training, as demonstrated by models trained on the re-scanned whole-slide Visium image (Supplementary Figure 7).

Much of the prior work on gene expression prediction from histology images has focused on modifying model architecture, exploring increasingly complex convolutional, graph, and transformer-based designs in an effort to improve predictive performance. Although these architectural modifications have produced incremental improvements, prediction accuracy remains highly variable across datasets, tissues, and technologies, indicating that model choice alone cannot fully explain observed performance differences. Consistent with this, replacing the ResNet50 feature extractor with the domain-adaptive foundation model UNI resulted in modest gains in PCC and reductions in normalized rMSE, while preserving the same relative performance trends across datasets (Supplementary Figure 3A-B). Similarly, applying RedeHist, a model specifically designed for gene expression prediction from histology images, showed the same relative performance trends, though with a slightly lower overall performance, likely because predicting gene expression at single-cell resolution is inherently more challenging than at patch-level resolution (Supplementary Figure 3C-D). Together, these results demonstrate that architectural modifications and foundation models can provide incremental gains, but they are ultimately constrained by the quality of the training data, emphasizing the need to prioritize data-centric approaches.

When predicting gene expression from histology images, accurately predicting cell-type marker genes is essential for determining cell-types and their spatial organization of tissues. Further, these genes are often more important for downstream biological interpretations. As such, performance for these genes may be more important than performance on other broadly expressed housekeeping genes for example. Using various cell-type marker gene-sets, we found nearly all marker genes had improved prediction performance when models were trained on Xenium data compared to Visium data in the breast cancer datasets (Supplementary Figure 11). These results indicate that higher-quality molecular training data improves the prediction of biologically informative, cell-type marker genes, reinforcing the importance of data quality considerations when developing and benchmarking spatial transcriptomics prediction models.

Although we found molecular data quality to impact prediction performance, other considerations may come into play in the curation of training data. For example, optimizing for molecular data quality generally comes at the expense of throughput. Therefore, if full-transcriptome gene expression prediction is prioritized, lower-quality datasets with higher throughput may be preferable for training. Cost constraints may also pose challenges to generating high-quality training data. Future work is needed to systematically investigate whether increasing the quantity of training data can compensate for reductions in data quality, and if so, to characterize the rate and extent of such trade-offs.

As with many deep learning applications, interpretability remains a key challenge. Although we can demonstrate quantifiable aspects of image data quality that lead to reduced prediction performance, how or why the models perform worse remains unclear. To this end, we applied Grad-CAM to demonstrate that even modest reductions in image resolution disrupted attention on the cellular and nuclear landmarks (Figure 4F), suggesting that cellular and nuclear landmarks may be important image features in prediction. We therefore hypothesized that predictions of cell-type-specific marker gene expression would direct model attention to corresponding cell-type features, analogous to a pathologist’s assessment. However, we found that even when we switched the predicted gene, attention remained fixed on general cellular and nuclear landmarks in a non-cell-type-specific manner (Supplementary Figure 12). This highlights the persistent challenges in interpreting deep learning interpretability methods. Going forward, the development of more robust explainability techniques, whether advances in Grad-CAM, SHAP [36], Integrated Gradients [37], or brand new approaches will be needed for deep-learning interpretability in this setting, though we anticipate such approaches will need to be paired with sufficiently high-quality images.

Overall, this study underscores the critical role of molecular and image data quality in spatial gene expression prediction from histology images. While recent developments have focused on improving modeling approaches, our findings highlight that improvements in training data quality offers an orthogonal strategy to tuning model architecture in enhancing predictive modeling using spatial transcriptomics. Our study further provides a comparative framework for evaluating future ST technologies to characterize aspects of molecular and image data quality to guide this evolving field. Looking forward, we anticipate a more holistic approach that integrates advancements in model architecture with improved generation and curation of training data is likely to yield the greatest performance gains in predictive modeling with spatial transcriptomics. Such a holistic approach will be important for not only improving prediction performance but also enhancing interpretability to ensure reliable and safe future applications in research and clinical settings.

## Methods

### Availability of data and materials

Two Xenium datasets (In Situ Replicate 1 and In Situ Replicate 2) along with an adjacent section of a Visium CytAssist dataset of a single breast cancer FFPE tissue block were obtained from the 10x Datasets website for *High-resolution mapping of the breast cancer tumor microenvironment using integrated single cell, spatial and in situ analysis of FFPE tissue* (https://www.10xgenomics.com/products/xenium-in-situ/preview-dataset-human-breast) [18]. Each dataset was accompanied by a H&Estained histology image. The Visium CytAssist dataset included three image resolutions: a low-resolution image (600 by 543 pixels), a high-resolution image (2000 by 1809 pixels), and a whole-slide image (20511 by 27587 pixels), with the high-resolution used in the primary analysis. The In Situ Replicate 1 and Replicate 2 Xenium datasets each included whole-slide histology images with resolutions of 20511 by 27587 pixels and 18728 by 27788 pixels, respectively. These images are herein referred to as whole-slide images (WSI). The Visium CytAssist dataset included spatial coordinates of the spots already aligned to its histology image, whereas the two Xenium datasets required aligning segmented cells to their corresponding histology images.

Three colon adenocarcinoma (COAD) datasets were obtained from the SPATCH web server (https://spatch.pku-genomics.org/) [27]. These datasets correspond to serial tissue sections assayed using Xenium 5K, VisiumHD, and CosMx 6K, respectively, and each is accompanied by a whole-slide H&E-stained histology image. The Xenium 5K, VisiumHD, and CosMx 6K whole-slide images have resolutions of 49221 by 36102 pixels, 44210 by 32542 pixels, and 46378 by 32812 pixels, respectively.

Source code for this study is available on GitHub at https://github.com/calebhallinan/dataquality_geneprediction.

### Restricting to shared tissue regions and shared genes

For the breast cancer datasets, we used STalign (v1.0.1) to bring the Visium and Xenium data to a common coordinate system by combining affine transformations with diffeomorphic metric mapping [19]. First, we aligned the Visium histology image (source) to the Xenium image (target) using an affine-only registration guided by eight manually selected landmarks. We then applied the learned affine transform to reposition each Visium spot onto the Xenium histology image. This procedure produced co-registered images of the same dimensions, allowing us to overlay Visium spot-level and Xenium single-cell data interchangeably on either histology image. Next, we registered the Xenium single-cell coordinates onto their provided histology image using STalign’s full pipeline: rasterize the mapped transcripts at 30*µ*m resolution, an initial affine alignment based on four manually placed landmarks followed by diffeomorphic metric mapping (parameters: *a* = 2500; *epV* = 1; *niter* = 2000; *sigmaA* = 0.11; *sigmaB* = 0.10; *sigmaM* = 0.15; *sigmaP* = 50; *muA* = [1,1,1]; *muB* = [0,0,0]; all other settings default). Finally, we identified the overlapping tissue region between the two technologies (Figure 1B). Likewise, we restricted our analyses to the 306 genes common to both assays, since Visium provides full-transcriptome coverage while Xenium targets a limited panel.

For the colon adenocarcinoma (COAD) datasets, each technology-specific dataset (Xenium 5K, VisiumHD, and CosMx 6K) were already aligned to its corresponding whole-slide histology image. Gene expression matrices were then restricted to genes shared across all three platforms, resulting in 2,506 retained genes for downstream analyses.

### Rasterization preprocessing to obtain common patch-resolution training data

After successfully aligning the breast cancer datasets, histology patches with associated gene expression data were created for input into the deep learning model. For the Visium dataset, each histology patch corresponded to exactly one Visium spot and its associated gene expression values. After aligning the Visium image with the Xenium image, each Visium spot corresponded to a patch size of ∼250x250 pixels. The center coordinates of these Visium patches were then used to generate corresponding patches on the Xenium image. Since the images were aligned, this approach ensured consistency in patch size and location across both datasets. Unlike Visium molecular data, Xenium molecular data corresponds to single cells, meaning each patch contained multiple cells. Therefore, the patch coordinates were used to aggregate (ie. pseudobulk) the Xenium gene expression data into a single spot, analogous to the Visium spots and as done by SEraster [20]. The resulting patch-resolution gene expression counts were then natural log-transformed with a pseudo-count of 1, and patches containing zero gene expression were excluded from both the Visium and Xenium datasets. The resulting histology patches, along with their associated patch-resolution gene expression matrices, were subsequently used as input for the deep learning model. There were a total of 3,958 patches for both datasets.

For the colon adenocarcinoma (COAD) datasets, histology patches with associated gene expression measurements were similarly generated to provide patch-resolution training data for the deep learning models. For each technology (Xenium 5K, VisiumHD, and CosMx 6K), spatial gene expression measurements were rasterized into a grid of non-overlapping square patches with size 55*µ*x55*µ* across the tissues. Gene expression values were aggregated within each patch by summing the raw counts from all cells for Xenium 5K and CosMx 6K and spots for VisiumHD that fell within the patch boundaries, thereby producing patch-level expression measurements that were directly comparable across technology despite differences in their spatial resolution. The resulting patch-resolution gene expression counts were natural log-transformed with a pseudo-count of 1, and patches with zero total gene expression were excluded from downstream analyses. The corresponding histology patches were extracted from the whole-slide images and paired with their aggregated gene expression matrices, giving a standardized patch-level dataset for model training and evaluation across all three COAD platforms.

### Gene imputation, sparsity, and noise data augmentation

Numerous gene imputation methods exist, each representing different functional classes [38]. For this analysis, we selected three methods: KNN [23], MAGIC [24], and SCVI [25].

KNN-smoothing smooths gene expression by identifying nearest neighbors based on partially smoothed and variance-stabilized expression profiles and aggregating their transcript counts. For this analysis, the number of neighbors was set to *k* =50, and *d* =20 top principal components were used to determine the nearest neighbors during each smoothing step. Patch 1587 within the dataset was excluded before applying KNN-smoothing and reintroduced afterward due to insufficient gene counts. The raw UMI counts were used as input, as recommended by the authors.

MAGIC (Markov Affinity-based Graph Imputation of Cells) smooths gene expression by employing a graph-based approach, where nodes represent individual cells and edges encode similarities between them, and diffusing gene expression information across this graph. For this analysis, the number of neighbors was set to *knn* = 50, and the diffusion operator’s power, *t*, was set to “auto.” All other parameters were kept at their default values. Log-transformed UMI counts with a pseudo-count of 1 were used as input, as recommended by the authors.

ScVI (Single-cell Variational Inference) leverages single-cell RNA-seq data to impute spatial transcriptomic gene expression using a neural network. The neural network was trained using the Chromium 5’ Gene Expression data from another serial section of the breast cancer dataset, along with 80% of the spatial data. All parameters were maintained at their default values, and the model was trained for a maximum of 300 epochs. Genes *TKT*, *SLC39A4*, and *GABARAPL2* were excluded because they were absent from the Chromium 5’ training dataset. Log-transformed UMI counts with a pseudo-count of 1 were used as input.

To introduce sparsity into the Xenium molecular data, Visium patches and their corresponding molecular data are used to impose sparsity on the aligned Xenium patches. Specifically, if a Visium patch contains counts *<*= *t* for gene *x*, the corresponding Xenium patch is set to zero for that gene. This procedure is applied across all patches and repeated at varying count thresholds *t* (0, 5, 10, 15, 20). This approach aims to closely replicate the sparsity observed in Visium molecular data within the Xenium dataset.

To introduce noise into the Xenium molecular data, for each patch we randomly sampled counts from a Poisson distribution with the expected value and variance of (*λ*) and added the sampled values to the original gene counts. The lambda (*λ*) parameter was varied from 5, 15, and 45 to simulate different noise levels.

### Neural network architectures and parameters

For image feature extraction, we used a pretrained ResNet50 model with the classification layer removed. The input consisted of extracted patches from histology images, sized 250x250x3 which were resized to the standard ResNet50 input size of 224x224x3. The patches were normalized per channel using the mean and standard deviation values from the pre-trained ResNet50 dataset. To enhance model robustness, the patches were subjected to random horizontal, vertical, and 90-degree rotations. The output of the ResNet50 model was the last layer of the model, a feature vector of size 2048, representing extracted image features. Once image features were extracted for each patch, we passed them through a four-layer multilayer perceptron (MLP) to both reduce dimensionality and learn the features most predictive of gene expression. Each MLP layer consisted of a linear transformation, batch normalization, ReLU activation, and 20% dropout. The final output is predicted gene expression for all 306 genes per patch. For training, mean-squared error (MSE) loss was used alongside the Adam optimizer [39], with a learning rate of 0.001 and weight decay of 0.00001. During training, a batch size of 64 was used, and each model was trained for 150 epochs. The datasets were divided into training (75%), validation (10%), and test (15%) sets. All results shown are on the test sets except the plotted predicted gene expression (Figure 2C) which is on the full dataset. Training was performed on an NVIDIA RTX 6000 Ada Generation GPU.

To evaluate the impact of an alternative feature extractor, we replaced the pretrained ResNet50 with the UNI domain-adaptive foundation model [21]. Histology patches were normalized and resized according to the UNI preprocessing protocol, and image features were extracted from the final embedding layer of the pretrained UNI model. These UNI-derived feature embeddings were then passed through the same four-layer MLP used in the ResNet50 model to predict patch-level gene expression. All other data augmentation strategies, training procedures, hyperparameters, dataset splits, and evaluation protocols were identical to those in the ResNet50-based models.

We additionally evaluated RedeHist, a deep learning model specifically designed for predicting gene expression from histology images at single-cell resolution [22]. RedeHist takes histology images, ST data, and scRNA-seq references as inputs to generate single-cell gene expression matrices with corresponding spatial coordinates and cell-type annotations. As the RedeHist GitHub repository provides a tutorial implementation for the breast cancer dataset, we followed this workflow to generate predictions for both the Visium and Xenium datasets using the associated scRNA-seq data as reference input (https://github.com/Roshan1992/RedeHist). Because RedeHist outputs gene expression at single-cell resolution, we aggregated the predicted expression values to patch resolution by constructing 250x250x3 image patches and summing gene expression within each patch, enabling direct comparison with the patch-level models.

### Deep learning interpretability method

To interpret the regions of the input image that the deep learning model attends to when predicting spatial gene expression, we employed Gradient-weighted Class Activation Mapping (Grad-CAM) [26]. Grad-CAM produces a coarse localization heatmap that highlights the most influential areas of the input image for a given gene prediction. This is achieved by computing the gradient of the target gene score with respect to the feature maps of the final convolutional layer within the ResNet50 architecture. These gradients are then globally averaged to obtain importance weights for each feature map. The weighted feature maps are subsequently combined through a linear summation, and a ReLU activation function is applied to retain only positively contributing regions. The resulting single-channel heatmap is up-sampled to match the original input image dimensions, yielding a visual representation of the regions the network considers most relevant for the given gene prediction.

### Overview of metrics used for evaluation

We measured the performance of predicting gene expression with the Pearson Correlation Coefficient (PCC) [40].

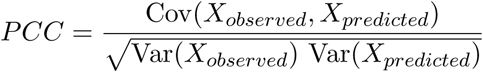

where

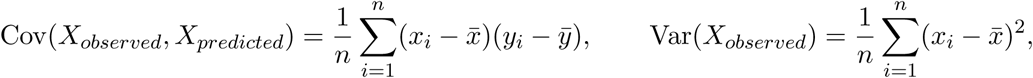

and similarly for Var(*X_predicted_*). *X_observed_* and *X_predicted_* are the observed (ground truth) and predicted gene expression, respectively. A higher PCC indicates better predictive performance.

We further measured prediction performance utilizing Normalized Root Mean Squared Error (rMSE), which quantifies the average magnitude of error between observed and predicted values, normalized by the range of observed expression values.

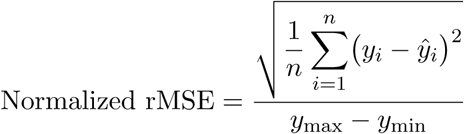

where *y_i_* denotes the observed (ground truth) gene expression for observation *i*, *ŷ_i_* denotes the corresponding model-predicted gene expression, and *y*_max_ and *y*_min_ are the maximum and minimum observed expression values, respectively. Lower normalized rMSE values indicate better predictive performance, with zero representing a perfect prediction.

## Acknowledgments

We thank JEFworks lab members Dee Velazquez, Sami Singh, Rafael dos Santos Peixoto, Manju Anant, and Kalen Clifton for their valuable feedback. We also thank Celia Hallinan for preparing the schematics in Figure 1 and for providing additional feedback on this work.

Research reported in this publication was supported by the National Institute Of General Medical Sciences of the National Institutes of Health under Award Number R35-GM142889 and the Smart Health and Biomedical Research in the Era of Artificial Intelligence and Advanced Data Science program under Award 2124230.

## Supplementary Figures

**Supplementary Figure 1:**
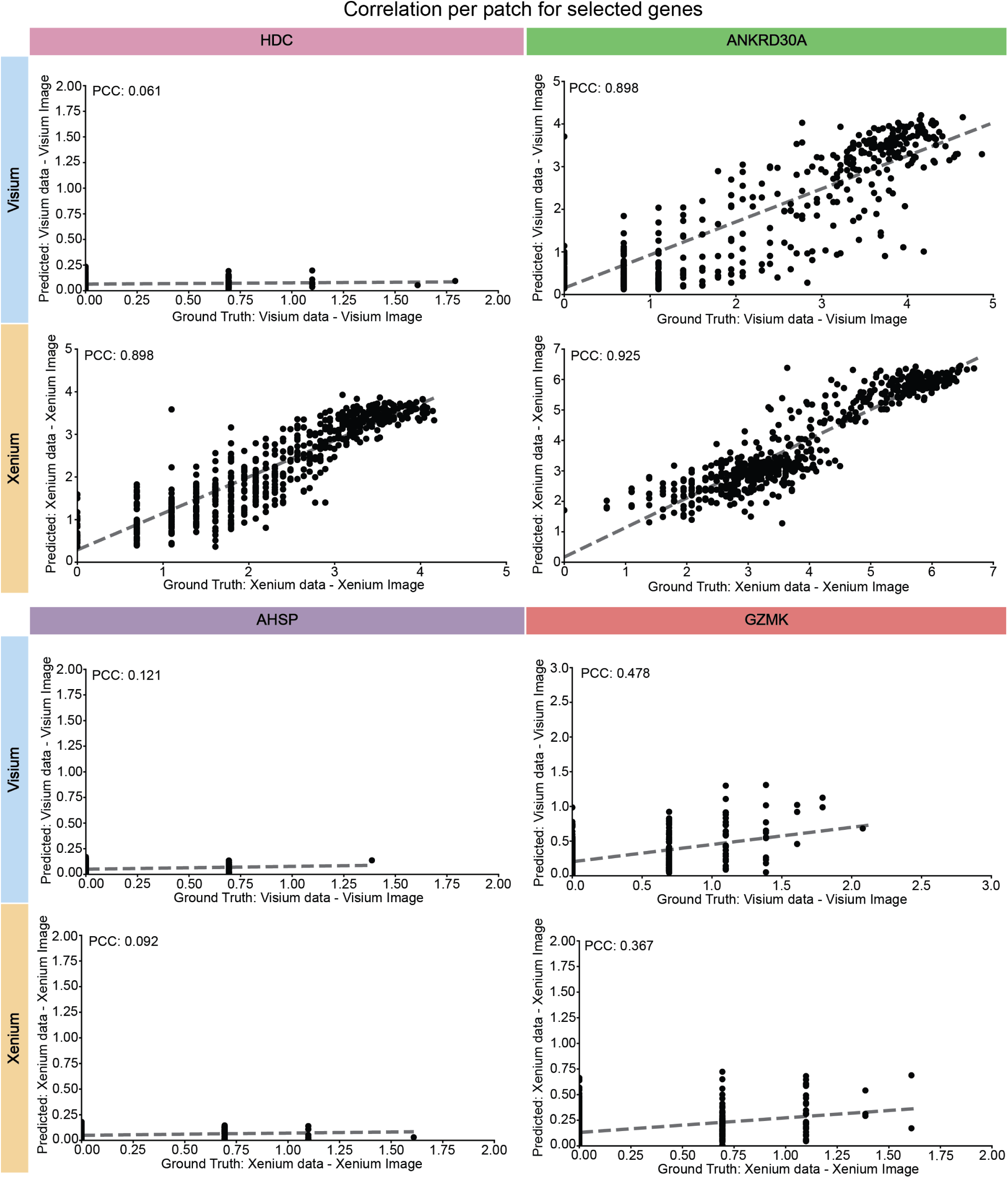
Correlation between ground-truth and predicted gene expression across patches for selected genes. Scatterplots comparing the ground truth held-out test set patches with the predicted expression of patches for *HDC*, *ANKRD30A*, *AHSP*, and *GZMK*. The gray dotted line denotes the linear regression fit. Analyses shown are based on a single seed rather than five independently trained models.

**Supplementary Figure 2:**
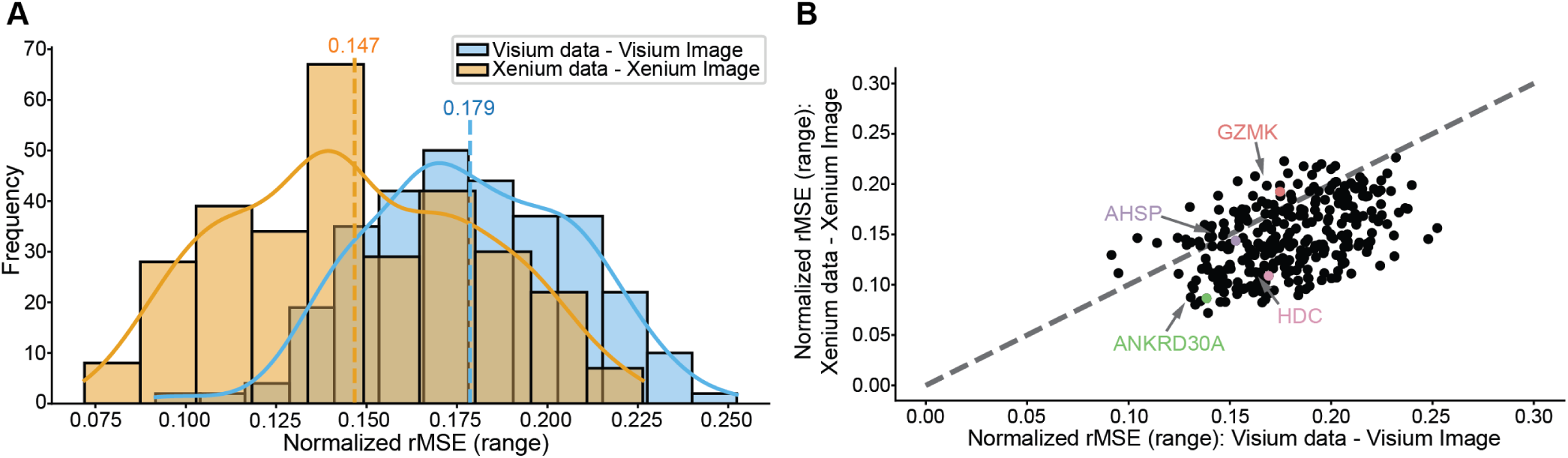
Prediction performance using normalized rMSE for Visium vs. Xenium data. **A.** Histogram showing the distribution of normalized rMSE for gene expression predictions using Visium and Xenium data. The dotted vertical line denotes the mean rMSE, and the solid curved line traces the density estimate. Results are computed on the test set and represent the average performance across five independently trained models. **B.** Scatterplot comparing the normalized rMSE of predictions from Visium and Xenium data, based on the test set and averaged over five models. The gray dotted line denotes x=y.

**Supplementary Figure 3:**
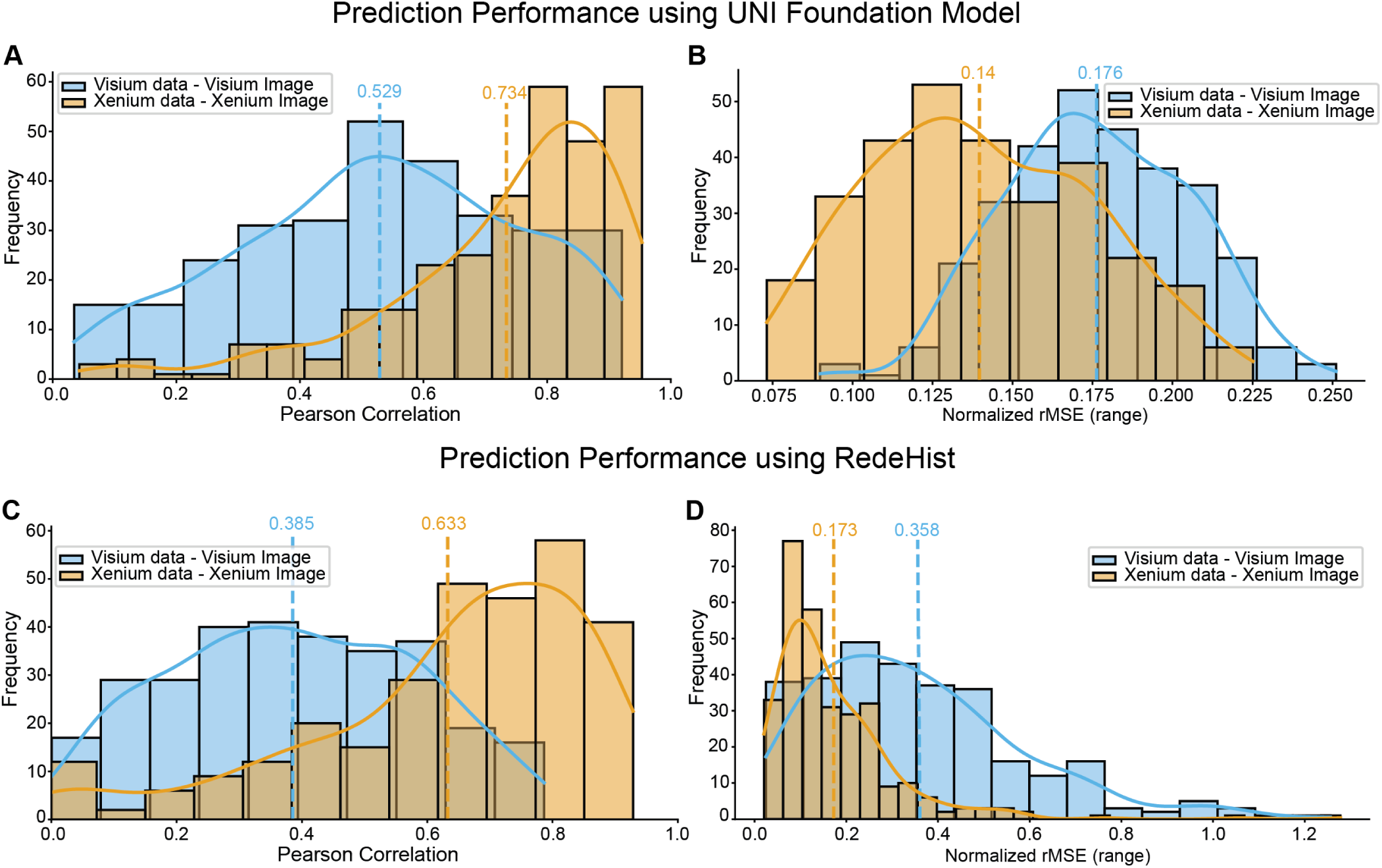
Prediction performance using the UNI foundation model and RedeHist. **A.** Histogram showing the distribution of Pearson correlation for gene expression predictions using Visium and Xenium data and trained using the UNI foundation model. The dotted vertical line denotes the mean Pearson Correlation, and the solid curved line traces the density estimate. Results are computed on the test set and represent the average performance across five independently trained models. **B.** Histogram showing the distribution of normalized rMSE for gene expression predictions using Visium and Xenium data and trained using the UNI foundation model. The dotted vertical line denotes the mean normalized rMSE, and the solid curved line traces the density estimate. Results are computed on the test set and represent the average performance across five independently trained models. **C.** Histogram showing the distribution of Pearson correlation for gene expression predictions using Visium and Xenium data and trained using RedeHist model. The dotted vertical line denotes the mean Pearson Correlation, and the solid curved line traces the density estimate. Results are computed on the test set and represent the average performance across five independently trained models. **D.** Histogram showing the distribution of normalized rMSE for gene expression predictions using Visium and Xenium data and trained using the RedeHist model. The dotted vertical line denotes the mean normalized rMSE, and the solid curved line traces the density estimate. Results are computed on the test set and represent the average performance across five independently trained models.

**Supplementary Figure 4:**
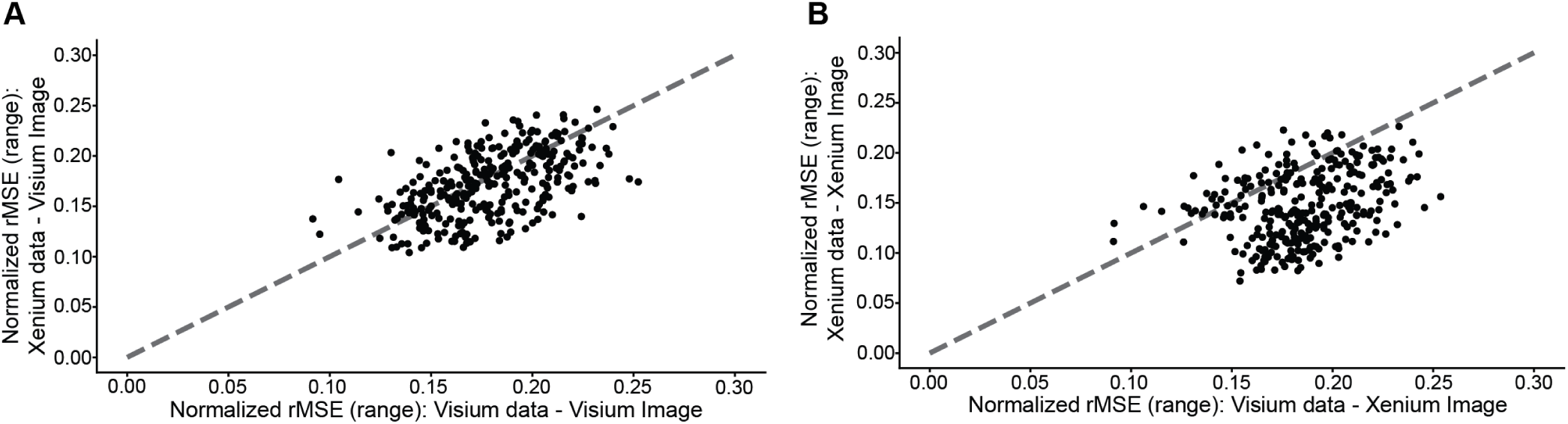
Prediction performance using normalized rMSE for Visium vs. Xenium molecular data. Scatterplots of normalized rMSE for models trained on varied molecular inputs, evaluated on the held-out test set and averaged across five independent runs, using (A) the Visium histology image and (B) the Xenium histology image. The gray dotted line denotes x=y.

**Supplementary Figure 5:**
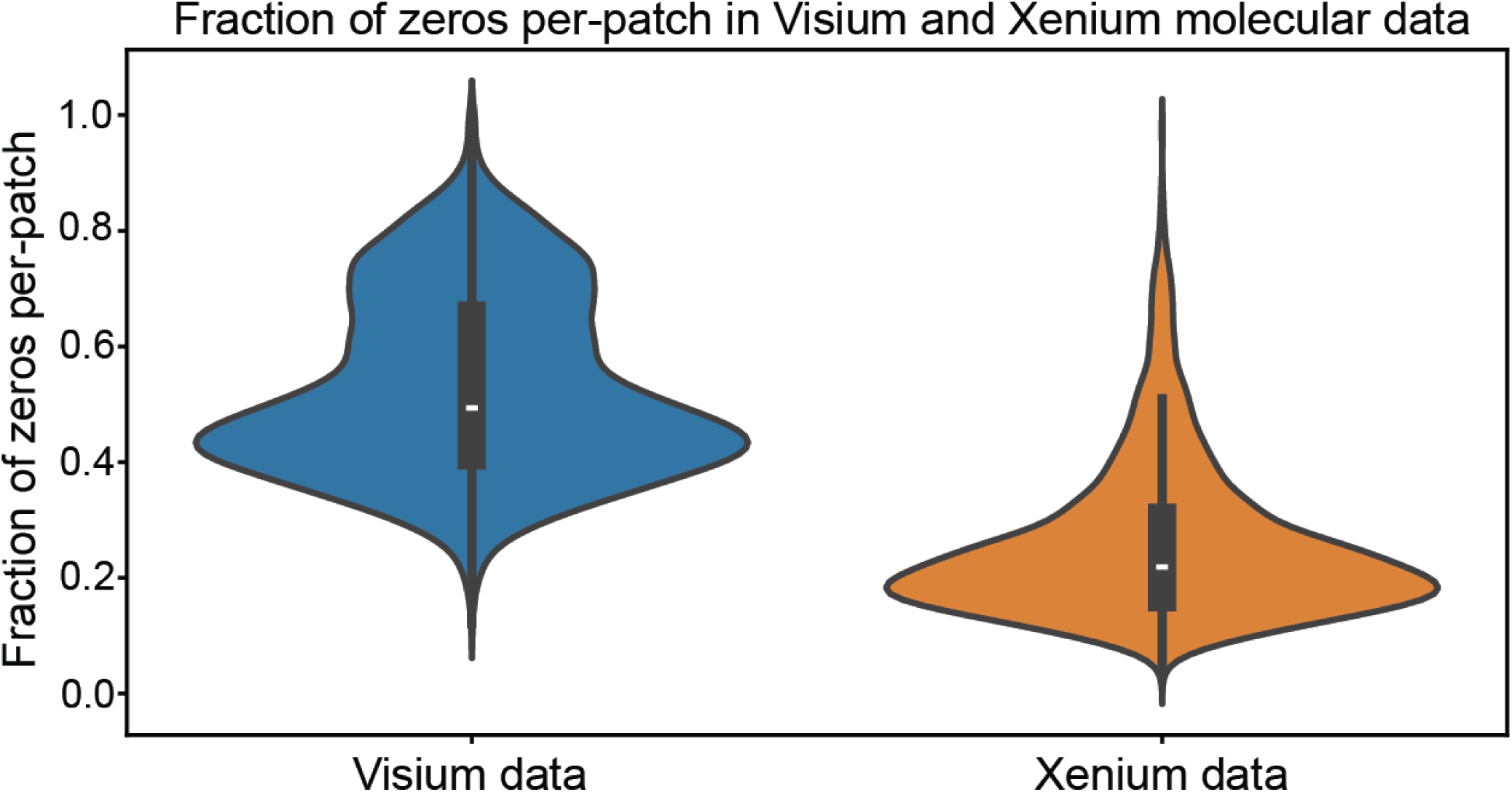
Sparsity in Visium and Xenium molecular data. Violin plots of the per-patch fraction of zero counts in Visium and Xenium molecular data. The shape of each violin reflects the density of values along the y-axis, and the overlaid boxplot indicates the median and the 25th and 75th percentiles.

**Supplementary Figure 6:**
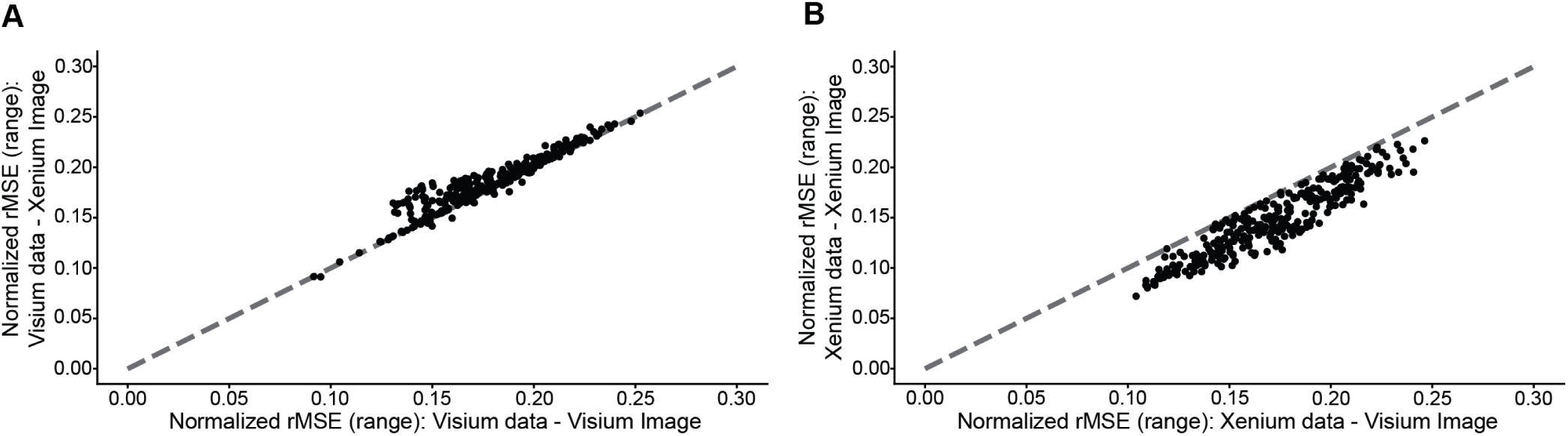
Prediction performance using normalized rMSE for Visium vs. Xenium imaging data. Scatterplots of normalized RMSE for models trained on varied image inputs, evaluated on the heldout test set and averaged across five independent runs, using (A) the Visium molecular data and (B) the Xenium molecular data. The gray dotted line denotes x=y.

**Supplementary Figure 7:**
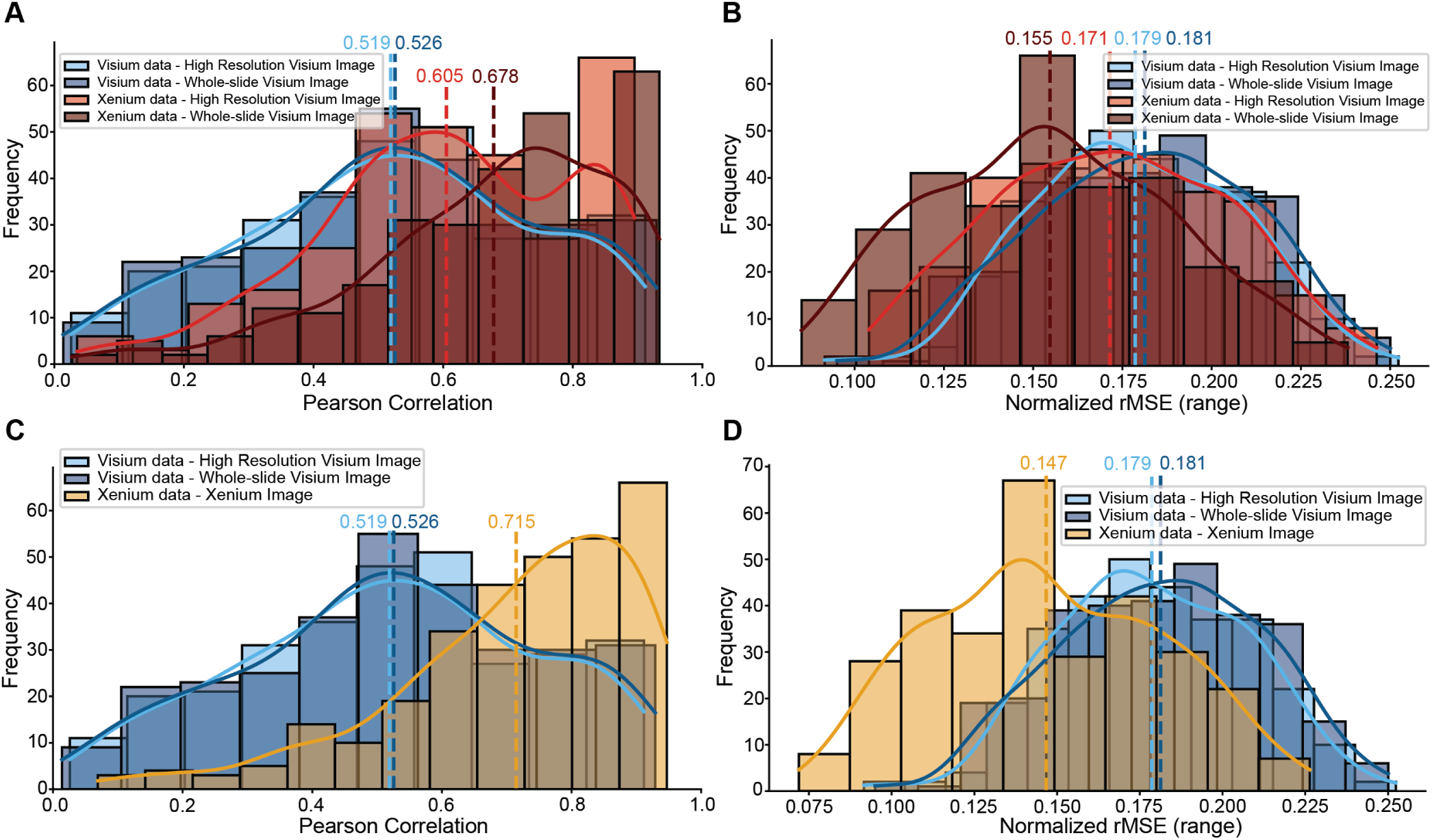
Prediction performance using the whole-slide Visium image. **A.** Histogram showing the distribution of the Pearson correlation for gene expression predictions using Visium molecular data with both the high-resolution Visium image and the Visium whole-slide image, as well as Xenium molecular data with both the high-resolution Visium image and the Visium whole-slide image. The dotted vertical line denotes the mean Pearson Correlation, and the solid curved line traces the density estimate. Results are computed on the test set and represent the average performance across five independently trained models. **B.** Histogram showing the distribution of the normalized rMSE for gene expression predictions using Visium molecular data with both the high-resolution Visium image and the Visium whole-slide image, as well as Xenium molecular data with both the high-resolution Visium image and the Visium whole-slide image. The dotted vertical line denotes the mean normalized rMSE, and the solid curved line traces the density estimate. Results are computed on the test set and represent the average performance across five independently trained models. **C.** Histogram showing the distribution of the Pearson correlation for gene expression predictions using Visium molecular data with both the high-resolution Visium image and the Visium whole-slide image, as well as Xenium molecular data with the Xenium image. The dotted vertical line denotes the mean Pearson Correlation, and the solid curved line traces the density estimate. Results are computed on the test set and represent the average performance across five independently trained models. **D.** Histogram showing the distribution of the normalized rMSE for gene expression predictions using Visium molecular data with both the high-resolution Visium image and the Visium whole-slide image, as well as Xenium molecular data with the Xenium image. The dotted vertical line denotes the mean normalized rMSE, and the solid curved line traces the density estimate. Results are computed on the test set and represent the average performance across five independently trained models.

**Supplementary Figure 8:**
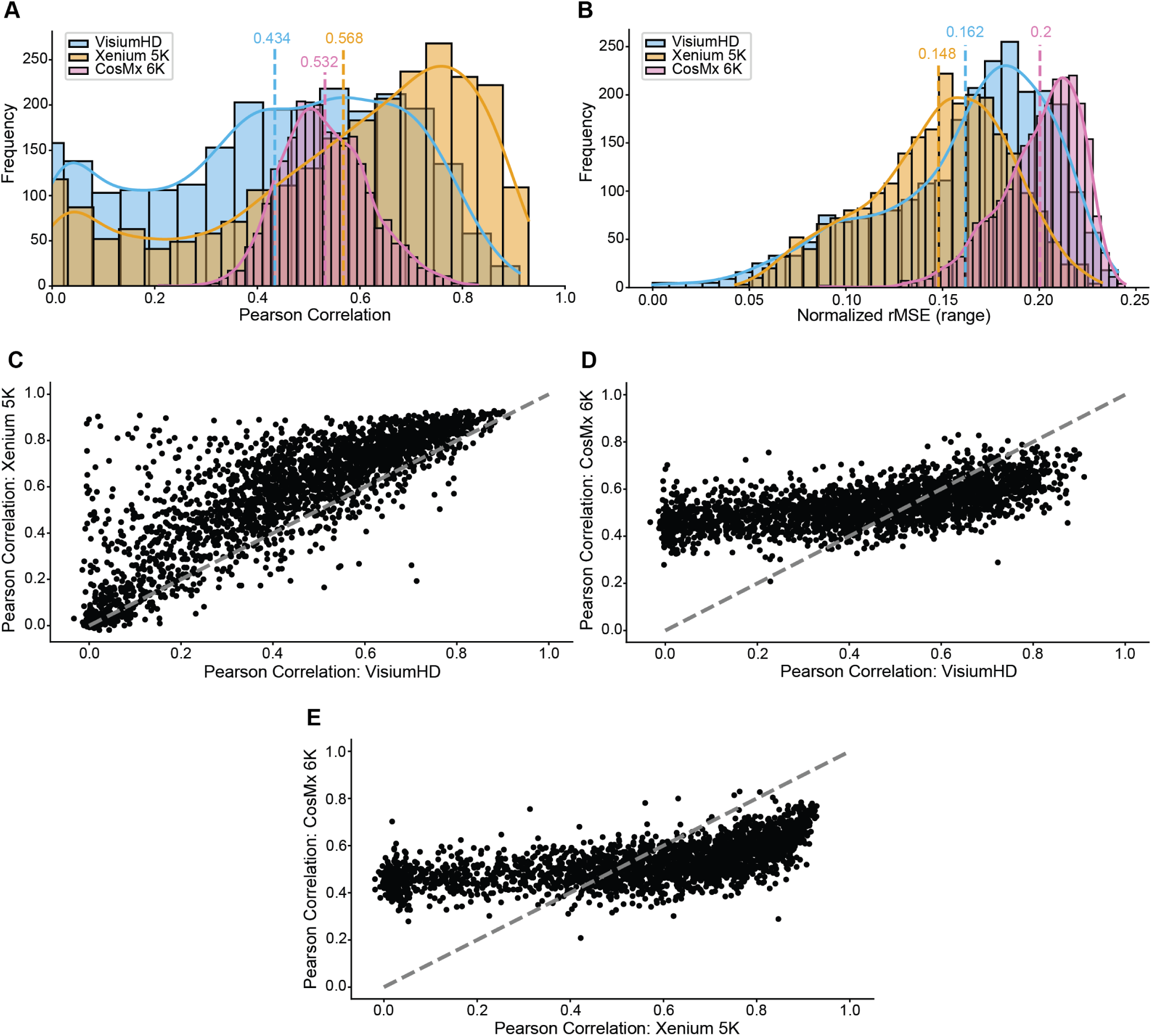
Prediction performance on the colon adenocarcinoma dataset. **A.** Histogram showing the distribution of Pearson correlation for gene expression predictions using VisiumHD, Xenium 5K, and CosMx 6K data. The dotted vertical line denotes the mean Pearson Correlation, and the solid curved line traces the density estimate. Results are computed on the test set and represent the average performance across five independently trained models. **B.** Histogram showing the distribution of normalized rMSE for gene expression predictions using VisiumHD, Xenium 5K, and CosMx 6K data. The dotted vertical line denotes the mean rMSE, and the solid curved line traces the density estimate. Results are computed on the test set and represent the average performance across five independently trained models. **C.** Scatterplot comparing the Pearson correlation of predictions from VisiumHD and Xenium 5K data, based on the test set and averaged over five models. The gray dotted line denotes x=y. **D.** Scatterplot comparing the Pearson correlation of predictions from VisiumHD and CosMx 6K data, based on the test set and averaged over five models. The gray dotted line denotes x=y. **E.** Scatterplot comparing the Pearson correlation of predictions from Xenium 5K and CosMx 6K data, based on the test set and averaged over five models. The gray dotted line denotes x=y.

**Supplementary Figure 9:**
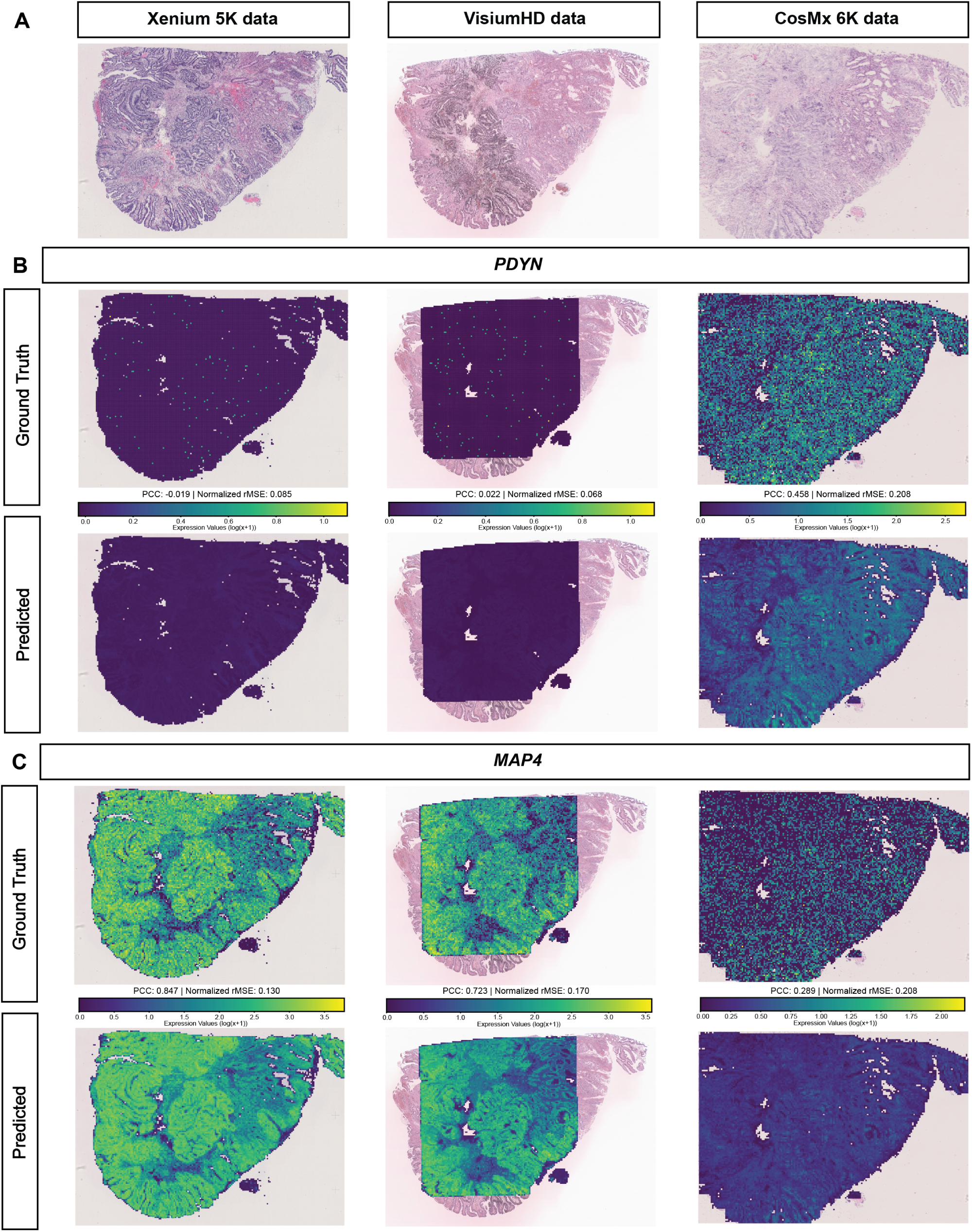
Examples of gene prediction performance on the colon adenocarcinoma dataset. **A.** Whole-slide H&E images associated with the Xenium 5K, VisiumHD, and CosMx 6K spatial transcriptomics technologies. **B.** Ground-truth and predicted spatial expression of the *PDYN* gene across Xenium 5K, VisiumHD, and CosMx 6K datasets. **C.** Ground-truth and predicted spatial expression of the *MAP4* gene across Xenium 5K, VisiumHD, and CosMx 6K datasets.

**Supplementary Figure 10:**
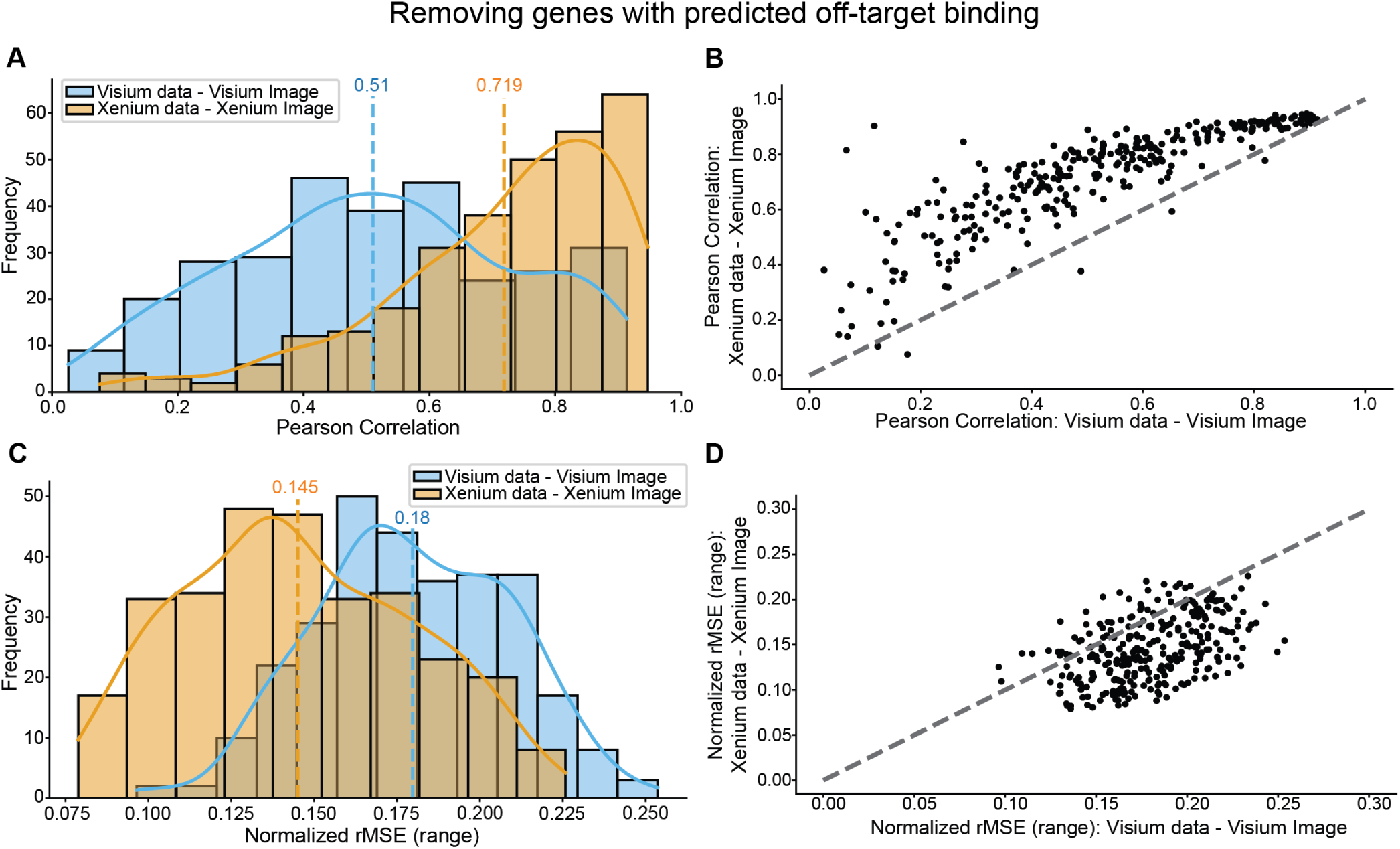
Visium vs. Xenium results when removing genes with predicted off-target binding. **A.** Histogram showing the distribution of Pearson correlation for gene expression predictions using Visium and Xenium data. The dotted vertical line denotes the mean Pearson Correlation, and the solid curved line traces the density estimate. Results are computed on the test set and represent the average performance across five independently trained models. **B.** Scatterplot comparing the Pearson correlation of predictions from Visium and Xenium data, based on the test set and averaged over five models. The gray dotted line denotes x=y. **C.** Histogram showing the distribution of normalized rMSE for gene expression predictions using Visium and Xenium data. The dotted vertical line denotes the mean rMSE, and the solid curved line traces the density estimate. Results are computed on the test set and represent the average performance across five independently trained models. **D.** Scatterplot comparing the normalized rMSE of predictions from Visium and Xenium data, based on the test set and averaged over five models. The gray dotted line denotes x=y.

**Supplementary Figure 11:**
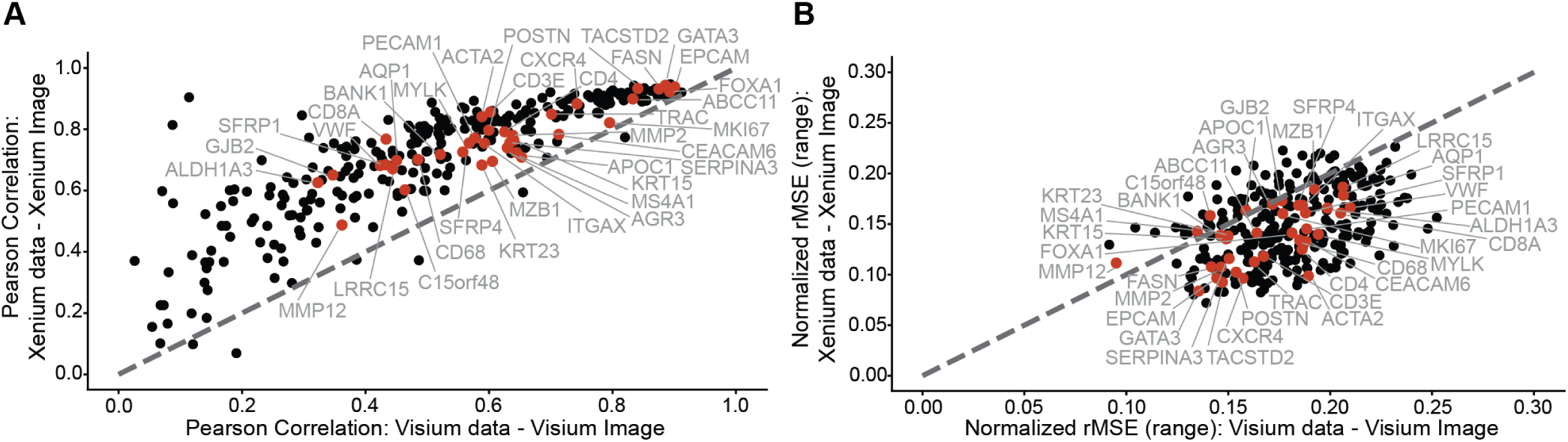
Cell-type marker gene prediction performance. **A.** Scatterplot comparing the Pearson correlation of predictions from Visium and Xenium data, based on the test set and averaged over five models. The gray dotted line denotes x=y, and cell-type marker genes are highlighted in red and labeled. **B.** Scatterplot comparing the normalized rMSE of predictions from Visium and Xenium data, based on the test set and averaged over five models. The gray dotted line denotes x=y, and cell-type marker genes are highlighted in red and labeled.

**Supplementary Figure 12:**
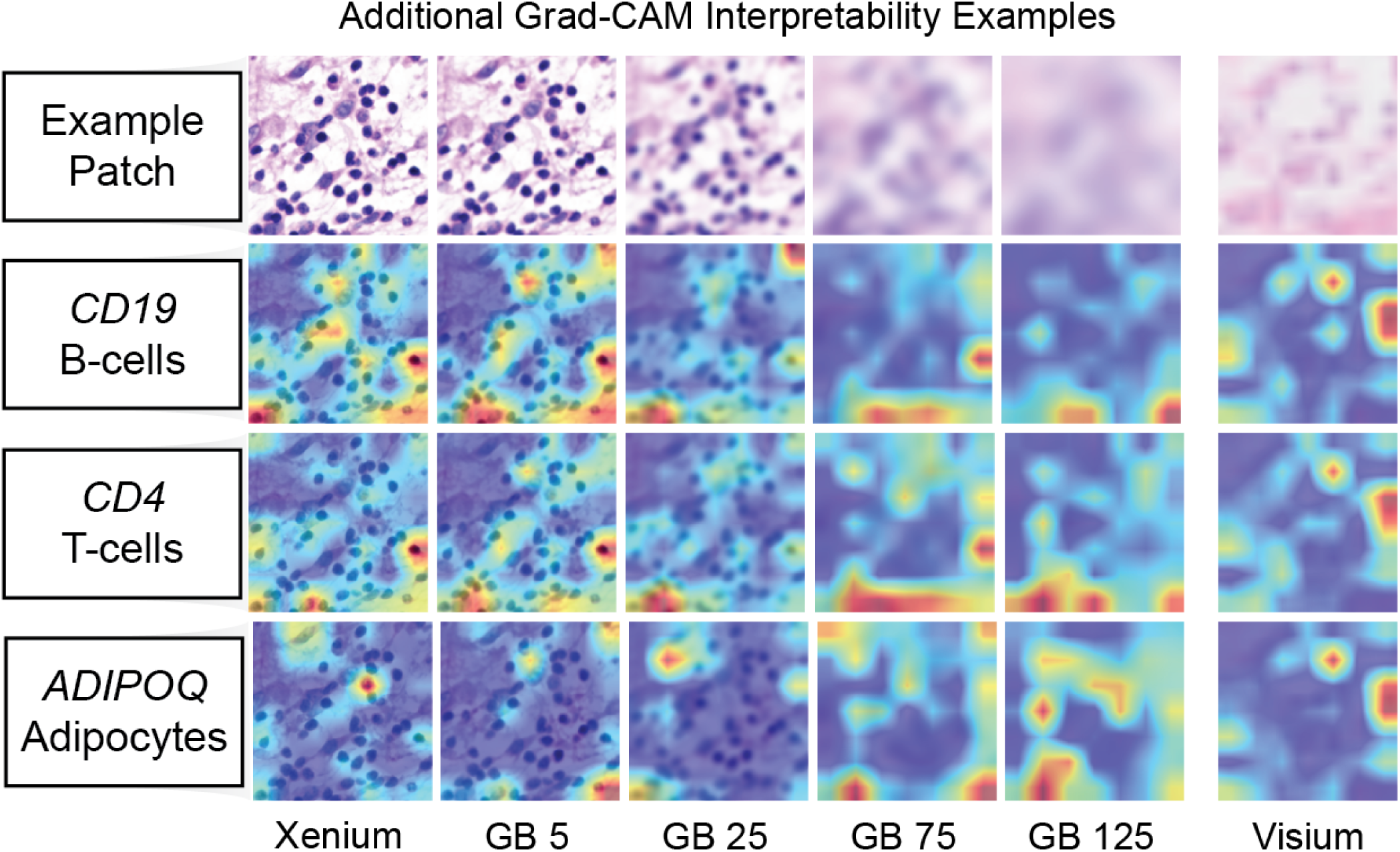
Additional Grad-CAM interpretability examples. A representative histology patch is shown under increasing Gaussian blur (GB) levels, with corresponding Grad-CAM heatmaps for *CD19* (B-cell marker), *CD4* (T-cell marker), and *ADIPOQ* (adipocyte marker).

